# Stem-like Prostate Remodeling in Obesity Mediates Resistance to 5α-Reductase Inhibition Therapy in BPH

**DOI:** 10.1101/2025.09.08.674969

**Authors:** Xingbo Long, Christina Sharkey, Rongbin Ge, Shuoshuo Wang, Yao Tang, Aaron Fleishman, Boqing Gu, Ruslan Korets, Boris Gershman, Heidi J. Rayala, Zongwei Wang, Aria F. Olumi

## Abstract

Benign prostatic hyperplasia (BPH) is closely associated with obesity and metabolic syndrome. Although 5α-reductase inhibitors (5ARIs) are widely used to treat BPH, their effectiveness varies significantly, particularly in obese patients. However, the mechanisms by which obesity affects prostate growth and modulates therapeutic responses remain poorly understood. In mice, high⍰fat diet (HFD) increased prostate weight and induced SRD5A2-independent epithelial remodeling with expansion of proximal urethral luminal epithelial cells. ScRNA-seq of prostate tissues from Srd5a2⍰null and control mice on HFD showed enrichment of proximal stem-like populations, reduced androgen signaling, and activation of WNT and NOTCH pathways via heightened stromal-epithelial crosstalk. Clinically, BMI gain, regardless of 5ARI use, correlated with transition zone growth, stronger activity of stem-cell signatures, reduced SRD5A2, and androgen signaling downregulation. Xenium spatial profiling indicated that BMI gain expands stem-like epithelial populations and a NOTCH⍰enriched periepithelial stromal niche surrounding epithelial compartments, supporting stemness. Patients with substantial BMI gain were less responsive to 5ARI, whereas weight control plus 5ARI therapy synergistically improved outcomes. In conclusion, BMI gain promotes proximal prostate enlargement through an SRD5A2-independent stem-like cell mediated pathway that blunts 5ARI efficacy. Combining pharmacotherapy with weight control yields superior efficacy and supports individualized management of BPH.

## Introduction

The global obesity epidemic has reached unprecedented levels across populations and imposing substantial socioeconomic burdens [1–3]. Beyond its cardiovascular and endocrine sequelae, obesity drives broad metabolic disturbances. Growing evidence now links benign prostatic hyperplasia (BPH), the most common chronic disorder affecting aging men, to obesity-related metabolic syndrome [4]. These observations support reframing BPH not merely as an age-dependent condition but as a metabolic disease fundamentally tied to the pathophysiological consequences of obesity.

Histologically, BPH is characterized by nonmalignant proliferation of epithelial and stromal components within the prostatic transition zone. Clinically, it is a leading cause of lower urinary tract symptoms (LUTS) in aging men and contributes substantially to diminished quality of life and healthcare costs [2]. Emerging data indicate that obesity, particularly visceral adiposity, is frequently comorbid with BPH and LUTS [5]. Several large prospective cohorts, including the Prostate Cancer Prevention Trial [6], the Osteoporotic Fractures in Men Study [7], and the Southern Community Cohort Study report that higher body mass index (BMI) is associated with an increased risk of LUTS [8, 9]. Observational studies further correlate higher BMI with larger prostate volume, elevated prostate⍰specific antigen (PSA) levels, and higher International Prostate Symptom Score (IPSS), reinforcing a role for metabolic dysregulation in BPH pathogenesis [10–12].

Steroid 5α-reductase type 2 (SRD5A2) underpins normal prostate development and BPH pathogenesis. Although 5ARIs, targeting SRD5A2⍰mediated dihydrotestosterone (DHT), are the only drugs that reduce prostate volume, responses are heterogeneous. We previously found that higher BMI and obesity⍰related inflammation are associated with SRD5A2 promoter hypermethylation and reduced prostatic expression [13]. Here we asked whether obesity promotes prostate growth through SRD5A2-dependent or -independent pathways that contribute to 5ARI resistance, and whether weight management could improve BPH outcomes and 5ARI efficacy [14, 15].

To address these questions, we integrated mouse single-cell RNA-seq with human Xenium spatial transcriptomics, bulk RNA-seq, and clinical cohort data to investigate how obesity influences prostate volume, cellular composition, *SRD5A2* expression, and responsiveness to 5ARI. Our analyses show that BMI gain promotes proximal prostate growth through an SRD5A2 independent stem-cell program that diminishes 5ARI efficacy. Integrating weight management with pharmacologic therapy improves outcomes, underscoring the need for personalized BPH treatment.

## Methods

### Mouse model generation and intervention

Detailed procedures for mouse model generation and experimental intervention have been described previously [16] and are provided in the Supplementary Methods.

### Clinical cohort

We obtained clinical information from patients enrolled in the MTOPS clinical trial [17]. This cohort consists of 3,047 patients, of whom 2,605 have intact baseline and follow-up data, with an average follow-up duration of 54.2 months. Clinical demographic characteristics of the MTOPS cohort included in this study are summarized in Supplementary Table 1. Baseline prostate transition biopsy samples from the MTOPS cohort were obtained from the National Institutes of Health/National Institute of Diabetes and Digestive and Kidney Diseases (NIH/NIDDK) Central Repository under an approved Resource Access Program (X01) application. Paired RNA-seq data from 108 baseline/follow-up biopsies from the MTOPS cohort were downloaded from Gene Expression Omnibus (GEO) (GSE262946) [18]. After excluding five patients with missing clinical data, 103 pairs were analyzed.

Prostate transurethral resection specimens from 33 patients were obtained from the biorepository at Beth Israel Deaconess Medical Center (BIDMC) for SRD5A2 immunohistochemical staining.

### Cohort information for Xenium analysis

All Xenium spatial transcriptomic data were derived from prostate tissue specimens collected from patients undergoing surgery for BPH at BIDMC between August 2020 and May 2023. A total of 63 patients and 142 tissue regions were included. The clinical characteristics of the patients, including age, BMI at surgery, preoperative BMI change, prostate size, and history of 5ARI, are summarized in Supplementary Table 2. Preoperative BMI change was defined as the difference between the most recent BMI value measured within six months to five years before surgery and the BMI value at the time of surgery. A positive value indicates an increase in BMI before surgery, while a negative value indicates a decrease. Total prostate volume was calculated following our previously published method [19]. Briefly, total prostate volume was measured on sagittal and axial T2-weighted MRI images using the prolate ellipsoid formula: Volume=0.52×length×width×height.

All clinical specimens obtained from BIDMC were collected with the approval of Institutional Review Board at BIDMC (IRB protocol #: 2019P000728, 2025P000620).

### Statistical analysis

All statistical analyses and graph generation were performed using R (version 4.4.3). Details regarding clinical parameter definition, clinical data analyses, bioinformatic processes and immunohistochemistry (IHC) are described in the Supplementary Methods.

## Results

### HFD drives prostate growth and epithelial remodeling independent of SRD5A2 expression

To delineate the molecular effects of obesity on prostate biology and evaluate the contribution of SRD5A2, *Srd5a2*-null (*Srd5a2^-/-^*) and *Srd5a2*-hetero (*Srd5a2*^+/-^) mice were subjected to either a RFD or HFD. Prostate tissues were subjected to histological analysis and scRNA-seq. Clinical relevance was assessed using Xenium spatial transcriptomics in our cohort and further validated in the MTOPS cohort (Figure 1A). HFD increased body weight and prostate weight across all genotypes (p < 0.05; Figure 1B, Left, Suppl Figure 1A), however, fold-change analysis demonstrated no significant differences in prostate weight between genotypes (Figure 1B, Right), suggesting SRD5A2 deletion does not affect HFD-induced prostate enlargement.

**Figure 1:**
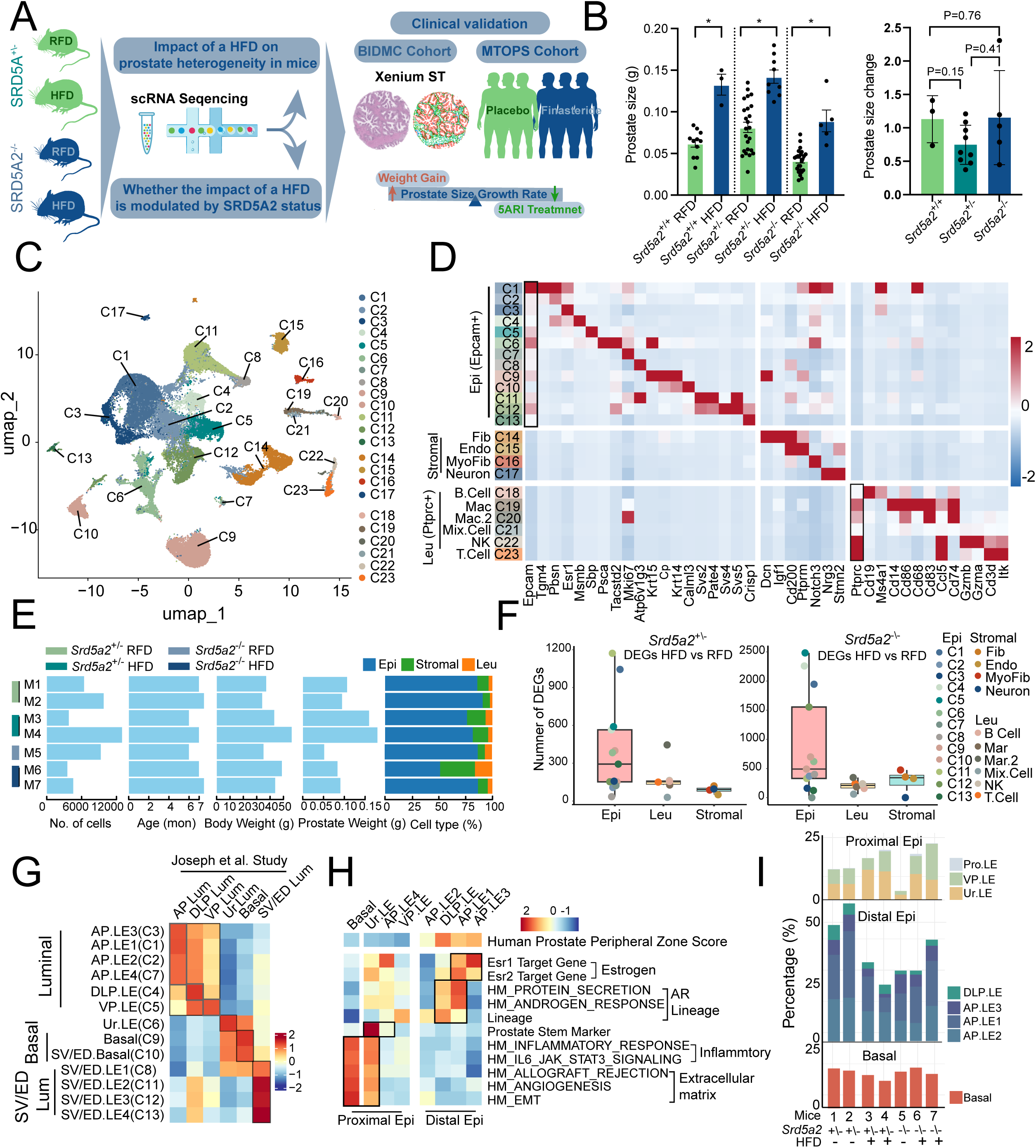
HFD drives prostate growth and epithelial remodeling independent of SRD5A2 expression. **A**: Experimental workflow to investigate the impact of high-fat diet (HFD) and SRD5A2 deficiency on prostate biology. **B**: Left: Bar graph showing absolute prostate weights in *Srd5a*2^-/-^ (knockout), *Srd5a2*^+/-^ (heterozygous) and *Srd5a2*^+/+^ (wild-type) mice under regular fat diet (RFD) and HFD conditions. Right: Fold-change in prostate weight (HFD vs. RFD) within each genotype. **C**: UMAP plot presenting the distribution of 23 cell clusters. **D**: Heatmap showing representative marker genes across 23 cell subtypes. **E**: Summary of sample metadata, sample grouping, age, body weight, prostate weight, cell counts, and major cell type distributions for the seven scRNA-seq prostate samples. **F**: Box plots display the number of differentially expressed genes (DEGs) between HFD- and RFD-treated mice in epithelial, leukocyte, and stromal compartments (*Srd5a2*^+/−^, left; *Srd5a2*^−/−^, right). **G**: SingleR correlation heatmaped epithelial clusters to established mouse prostate epithelial subtypes from the Joseph et al. database [14]. Clusters were annotated by anatomical identity: anterior prostate (AP), dorsolateral prostate (DLP), ventral prostate (VP), urethral prostate (Ur), basal, and seminal vesicle/ejaculatory duct (SV/ED). **H**: Heatmap displays Gene Set Variation Analysis (GSVA) enrichment scores of different gene sets across epithelial clusters (The source of gene sets were summarized in Suppl Table S3). HM: hallmark. **I**: Bar plots showing the proportion of epithelial cells from different anatomical region in each scRNA sample. Fib: Fibroblast; Endo: Endothelial cell; Myofib: Myofibroblast; Mac: Macrophage; NK: Natural killer cell. Epi: Epithelium, Leu: Leukocyte.

To map HFD-induced cellular heterogeneity, we performed scRNA-seq on prostate tissues from *Srd5a2^-/-^* and *Srd5a2*^+/-^ mice (n=7), identifying 23 cell clusters (Figure 1C). Clusters were annotated based on canonical markers: epithelial cells (C1-C13), stromal cells (fibroblasts C14, endothelial C15, myofibroblasts C16, neurons C17), immune cells (B cells C18, NK cells C22, T cells C23, macrophage subsets C19-C20), and a mixed population with multilineage markers (C21) (Figure 1D). Figure 1E summarizes biological information, cell counts and major cell type distributions across the seven samples.

To assess the effect of HFD on transcription across prostate compartments, we quantified differentially expressed genes (DEGs) between HFD- and RFD-treated mice in epithelial cells, leukocytes, and stromal cells under *Srd5a2*^+/-^ and *Srd5a2*^-/-^ backgrounds. As shown in Figure 1F, epithelial compartments exhibited significantly more DEGs and greater transcriptional heterogeneity than other compartments, regardless of SRD5A2 status. These findings establish the epithelium as the primary dietary stress-responsive population and thus the focus for subsequent analyses.

To further annotate the murine epithelial clusters and define their spatial identity, we used a previously characterized framework that maps murine epithelial populations to anatomical subregions of the mouse prostate epithelium [20] (Figure 1G). Region-specific clusters, including anterior prostate (AP), dorsolateral prostate (DLP), ventral prostate (VP), urethral (Ur) luminal cells (LE), basal, and seminal vesicle/ejaculatory duct (SV/ED), formed distinct groups with characteristic transcriptional signatures. Based on their anatomical locations, we annotated these epithelial clusters accordingly (Figure 1G). Non-prostatic SV/ED clusters were excluded as potential dissection artifacts. To gain further insight into the functional heterogeneity of epithelial subpopulations, we evaluated gene signature enrichment across epithelial clusters (Figure 1H). We first applied a set of luminal cell-specific genes from the human prostate peripheral zone (PZ) to map mouse epithelial clusters to spatial regions of the human prostate epithelium (Suppl Table 3) [21]. Mouse epithelial clusters AP.LE-3 and DLP.LE exhibited higher PZ genes enrichment, indicating transcriptional similarity to distal human epithelial compartments of the PZ. Conversely, Basal, Ur.LE, AP.LE4, and VP.LE were transcriptionally similar to proximal regions of the human prostate (Figure 1H, Suppl Figure 1B).

Functionally, proximal cells (particularly Ur.LE) were enriched for stemness and inflammation pathways, while distal cells exhibited hormone-regulated heterogeneity, including estrogen-dominant AP.LE3, AR-enriched DLP.LE, and bipotent AP.LE1 subtypes. Notably, HFD induced a marked expansion of proximal epithelial cells, especially the Ur.LE population, in both *Srd5a2*^+/-^ and *Srd5a2-/-* mice prostate (Figure 1I).

### HFD promotes proximal prostate expansion and stem-like reprogramming via SRD5A2-independent mechanisms

To explore the developmental trajectories of prostate epithelial cells, we conducted pseudotime analysis using Monocle 2. Pseudotime analysis revealed a stem-like origin in the urethral/proximal epithelium (Component 1), followed by a branchpoint that bifurcated into two distinct differentiation trajectories (Figure 2A-B). The first trajectory (Component 2) represented an androgen-responsive (AR) lineage, culminating in AR-driven DLP.LE cells. The second trajectory (Component 2) followed an estrogen-responsive pathway, progressing from bipotent AP.LE1 cells to estrogen-driven AP.LE3 cells (Figure 2B). Signature analysis confirmed that AR activity increased along Component 2, while estrogen receptor (ESR1) target gene expression inversely correlated with Component 2. Stem-like gene signatures peaked near the pseudotime origin and showed strong negative correlation with Component 1, supporting the urethral/proximal region as a progenitor-rich niche (Figure 2A and C). Furthermore, we found that HFD when compared to RFD significantly altered murine epithelial cell distribution across prostate regions, increasing urethral/proximal populations (47.6% vs 27.7%) while decreasing distal representation (52.33% vs 72.14%) (Figure 2D). Our results demonstrate that HFD preferentially drives the expansion of urethral/proximal epithelial populations while suppressing distal differentiation. Importantly, this urethral/proximal enrichment pattern persisted in *Srd5a2*^-/-^ mice despite the deletion of SRD5A2 (Suppl Figure 2A).

**Figure 2:**
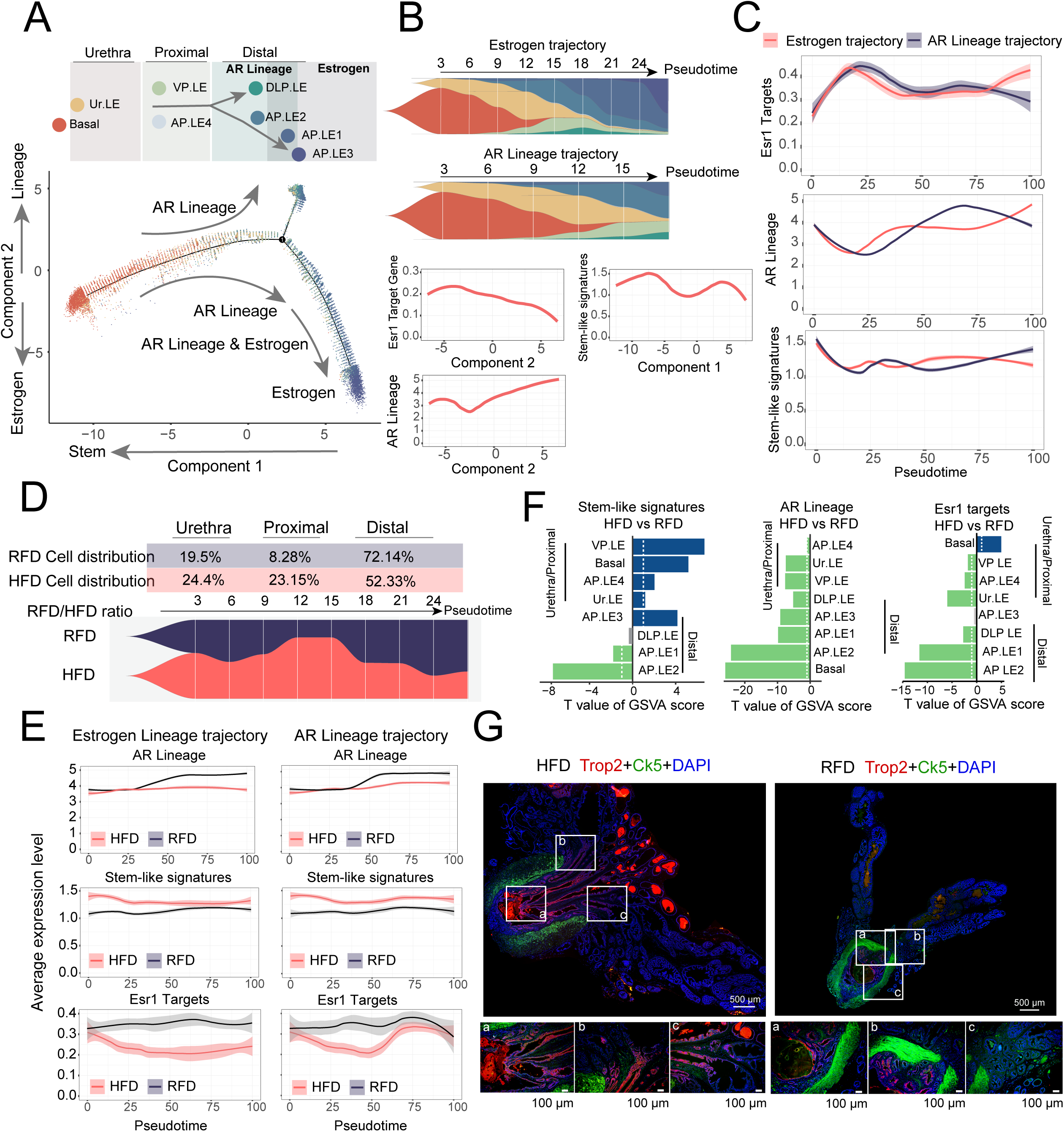
Epithelial differentiation trajectories and HFD-induced changes. **A**: Top: Schematic summarize of epithelial trajectory patterns inferred from pseudotime and cell distribution. Trajectories are grouped by anatomical origin of urethral (Ur.LE), proximal (VP.LE, AP.LE4), and distal (DLP.LE, AP.LE1-3) regions. Bottom: Monocle 2 inferred pseudotime trajectory of epithelial cells branching into AR lineage and estrogen trajectories. Each dot represents a single cell, colored by cluster (all samples). **B**: Top: Fishplot showing epithelial cell distribution across pseudotime. Bottom: LOESS-fitted average expression of ESR1 target, AR lineage, and stem-like gene signatures along the two Monocle components. **C**: LOESS curves showing ESR1 target, AR lineage, and stem-like gene signatures expression across pseudotime in estrogen and AR lineage trajectories. **D**: Fishplot showing pseudotime distribution of epithelial cells under HFD vs RFD in *Srd5a2*^+/-^ mice. **E**: LOESS-fitted average expression of ESR1 target, AR lineage, and stem-like gene signatures across pseudotime in estrogen and AR trajectories under *Srd5a2*^+/-^ condition, comparing HFD and RFD groups. **F**: Gene Set Variation Analysis (GSVA) based comparison of pathway activity scores between HFD and RFD across epithelial cell types in *Srd5a2*^+/-^ mice, shown as t-values from a linear model. **G**: Immunofluorescence of Ur.LE marker Trop2 and basal cell marker CK5 in prostates of *Srd5a2*^+/-^ mice under HFD and RFD.

Beyond altering cellular distribution patterns, HFD induced spatial reprogramming of steroid hormone and stemness signatures, enhancing stem-related genes in proximal regions while suppressing steroid-responsive genes distally across both trajectories (Figure 2E and F, Suppl Figure 2B). Notably, although *Srd5a2*^-/-^ mice exhibited attenuated androgenic and estrogenic signaling, proximal enrichment of stemness signatures remained robust (Suppl Figure 2C, D and E). This was corroborated by immunofluorescence, which demonstrated HFD-specific expansion of Trop2^+^ stem-like Ur.LE cells in urethral/proximal prostate regions (Figure 2G). Quantification of Trop2+ area as a percentage of total prostate confirmed significant HFD-induced expansion of the urethral/proximal region, a phenotype that persisted despite the absence of SRD5A2 in the *Srd5a2*^-/-^ mice (Suppl Figure 2F-H).

### HFD promotes stromal-epithelial crosstalk in proximal prostate

Emerging evidence indicates that stromal-epithelial crosstalk plays a crucial role in regulating prostate epithelial cell growth [22]. To elucidate the mechanisms driving HFD-induced proximal prostate enlargement, we performed ligand-receptor analysis. As shown in Figure 3A, myofibroblasts and fibroblasts are key ligand signaling sources (“senders”), with Ur.LE and basal cells are the primary signal “receivers”. Furthermore, the proximal region of the prostate receives more stromal-derived signaling input (Figure 3B), suggesting the proximal epithelium has enhanced signaling-receiving capacity compared to the distal epithelium, which relies mainly on steroid hormones. Moreover, HFD strengthened the stromal-Ur.LE and proximal stromal-epithelial interactions (Figure 3B and C).

**Figure 3:**
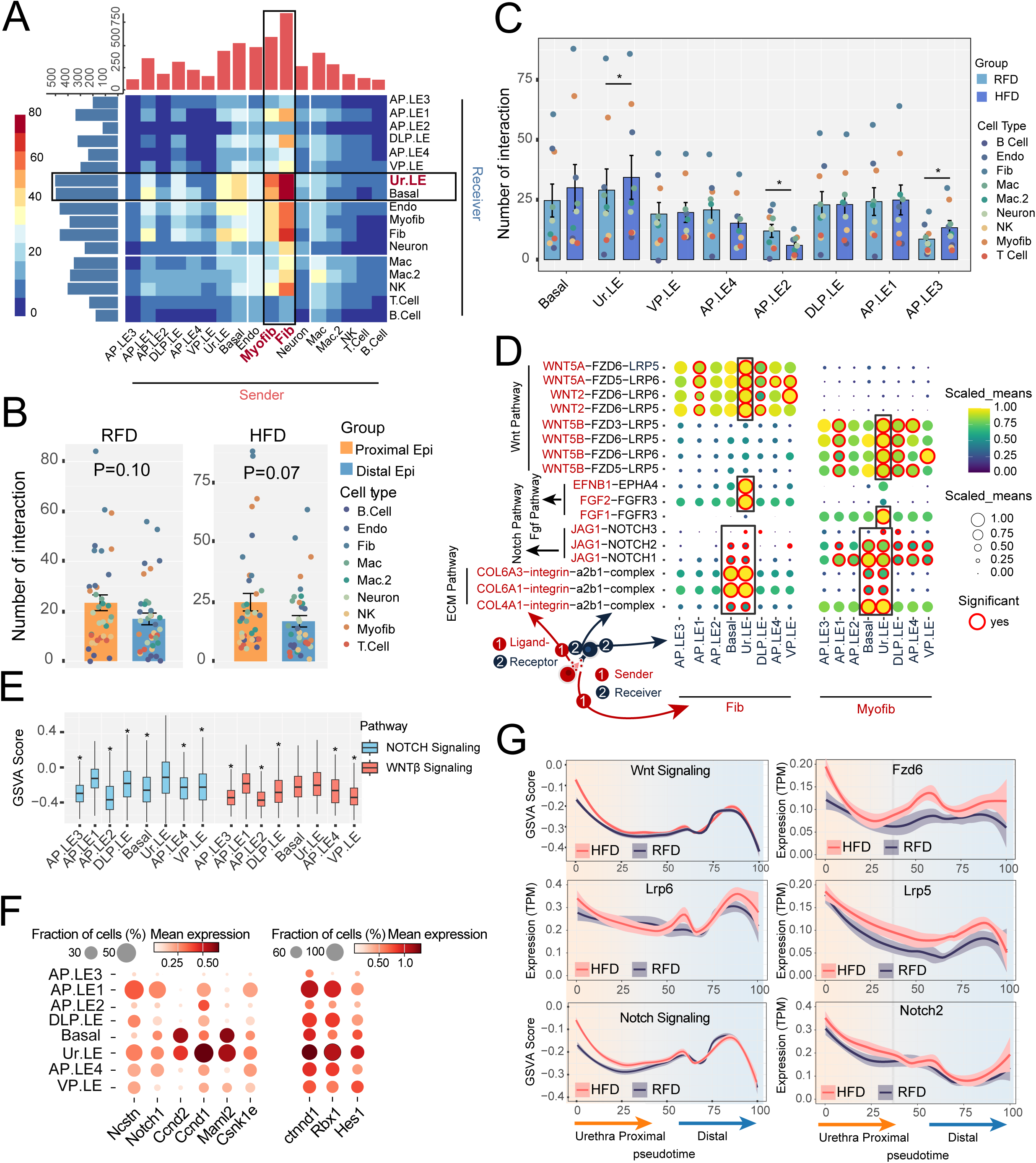
HFD promotes stromal-epithelial crosstalk in proximal prostate. **A**: Heatmap showing the number of predicted ligand-receptor pairs predicted by CellPhoneDB between cell types. Bar plots indicate the total number of ligand-receptor senders and receivers per cell type. **B**: Bar plots showing differences in the number of stromal-epithelial interactions received by proximal and distal epithelial cells under HFD and RFD. **C**: Bar plots comparing the number of stromal-derived ligand-receptor pairs received by epithelial cell types under HFD vs RFD. *P < 0.05. **D**: Dot plot showing epithelial cell-specific ligand-receptor interactions involving Wnt, Notch, and related pathways from fibroblasts and myofibroblasts. **E**: Gene Set Variation Analysis (GSVA) analysis showing Wnt and Notch pathway enrichment (Hallmark gene set) across epithelial cell subtypes. *Ur.LE significantly enriched vs other epithelial subtypes (P < 0.05). **F**: Dot plot showing expression of representative Wnt and Notch pathway genes across epithelial cells. Data merged from all samples for figure A to F. **G**: LOESS-smoothed GSVA scores of Wnt/Notch pathways and receptor expression (TPM: Transcripts Per Million) along pseudotime in Srd5a2^+/-^ mice under HFD and RFD. Fib: Fibroblast; Endo: Endothelial cell; Myofib: Myofibroblast; Mac: Macrophage; NK: Natural killer cell.

In addition, we found that androgen signaling interaction, mediated by key enzymes SRD5A2 and HSD17B12 originates from fibroblasts, macrophages, and neurons, while estrogen signaling interaction comes from endothelial cells and macrophages (Suppl Figure 3A). In *Srd5a2*^+/-^ mice, HFD reduced both androgen/estrogen interactions (Suppl Figure 3A) and SRD5A2 expression but had minimal effect on HSD17B12 expression (Suppl Figure 3B and C). In *Srd5a2*^-/-^ mice, androgen signaling interactions were nearly abolished (Suppl Figure 3A and B), whereas estrogen signaling interactions were markedly enhanced and remained unaffected by HFD (Suppl Figure 3A).

Lastly, we found that compared to other epithelial cells, urethral Ur.LE/basal epithelial cells exhibited stronger interactions with stromal cells through multiple growth- and stemness-associated pathways, such as WNT, FGF, and NOTCH (Figure 3D). Consistently, Enrichment analysis confirmed WNT and NOTCH pathways and their related genes were enriched in Ur.LE (Figure 3E and F). Notably, WNT/NOTCH pathway enrichment in proximal prostate was further amplified by HFD through increased stromal-derived interactions (Suppl Figure 4A) and upregulated expression of WNT/NOTCH receptors (Fzd6, Lrp5/6, Notch2) in proximal epithelial cells, independent of Srd5a2 status (Figure 3G and Suppl Figure 4B).

### Body weight gain is associated with total and transition zone prostate growth

To strengthen the translational relevance of our murine findings, we analyzed MTOPS clinical data to correlate BMI change with zonal prostate growth rates. Multivariable analysis revealed that patients who received 5ARI exhibited significantly reduced prostate growth rates when assessing volumes of total prostate (βx100=-29.86, 95% CI: -34.16 to -25.57), transition zone (βx100=-32.6, 95% CI: -39.51 to -25.70) and peripheral zone (βx100=-24.1, 95% CI: -36.51 to - 11.79) (all p<0.001; Figure 4A-C). Notably, longitudinal BMI change predicted the growths of total prostate volume (βx100=0.99, 95% CI:0.06 to 1.92) and transition zone volume (βx100=1.67, 95% CI:0.17 to 3.16), but not the growths of peripheral zone volume (Figure 4A-C). Univariable analysis demonstrated similar results (Suppl Figure 5). These clinical observations align with our mouse findings of HFD-driven expansion of prostate proximal region.

**Figure 4:**
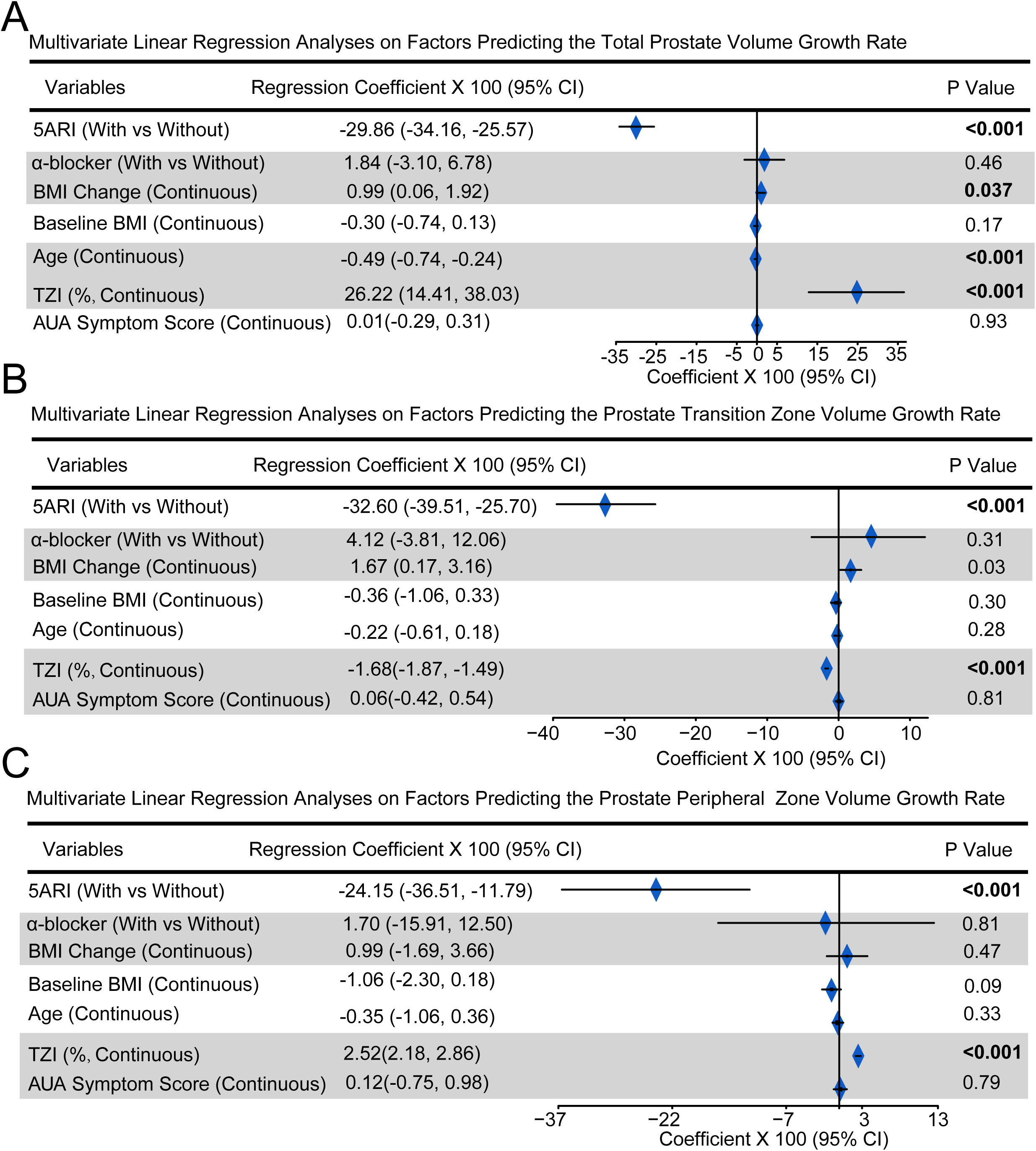
Body weight gain is associated with the growth of prostate transition zone. **A-C**: Multivariate linear regression analyses on factors predicting the growth rate of total prostate volume (A), prostate transition zone volume (B) and prostate peripheral zone volume (C).

### Body weight gain expands stem-like epithelium via NOTCH-enriched niches

To validate the murine finding that HFD enhances proximal prostate stemness in humans, we performed high-resolution Xenium spatial transcriptomic profiling of 142 BPH tissue regions (Suppl Figure 6A and 6B, Table S4). BMI gain was associated with expansion of stem-like club cells (Ur.LE homologs) and increased prostate volume, accompanied by reduced luminal cells and diminished SRD5A2 expression(Figure 5A and B). These associations were independent of 5ARI use (Suppl Figure 6C). IHC confirmed that BMI gain was inversely correlated with SRD5A2 expression (Suppl Figure 6D). Consistently, MTOPS RNA-seq data demonstrated that longitudinal BMI gain was accompanied by enrichment of stem-like club/hillock programs and a concurrent reduction in AR/luminal signatures and serum DHT levels, regardless of 5ARI treatment (Suppl Figure 6E–G).

**Figure 5:**
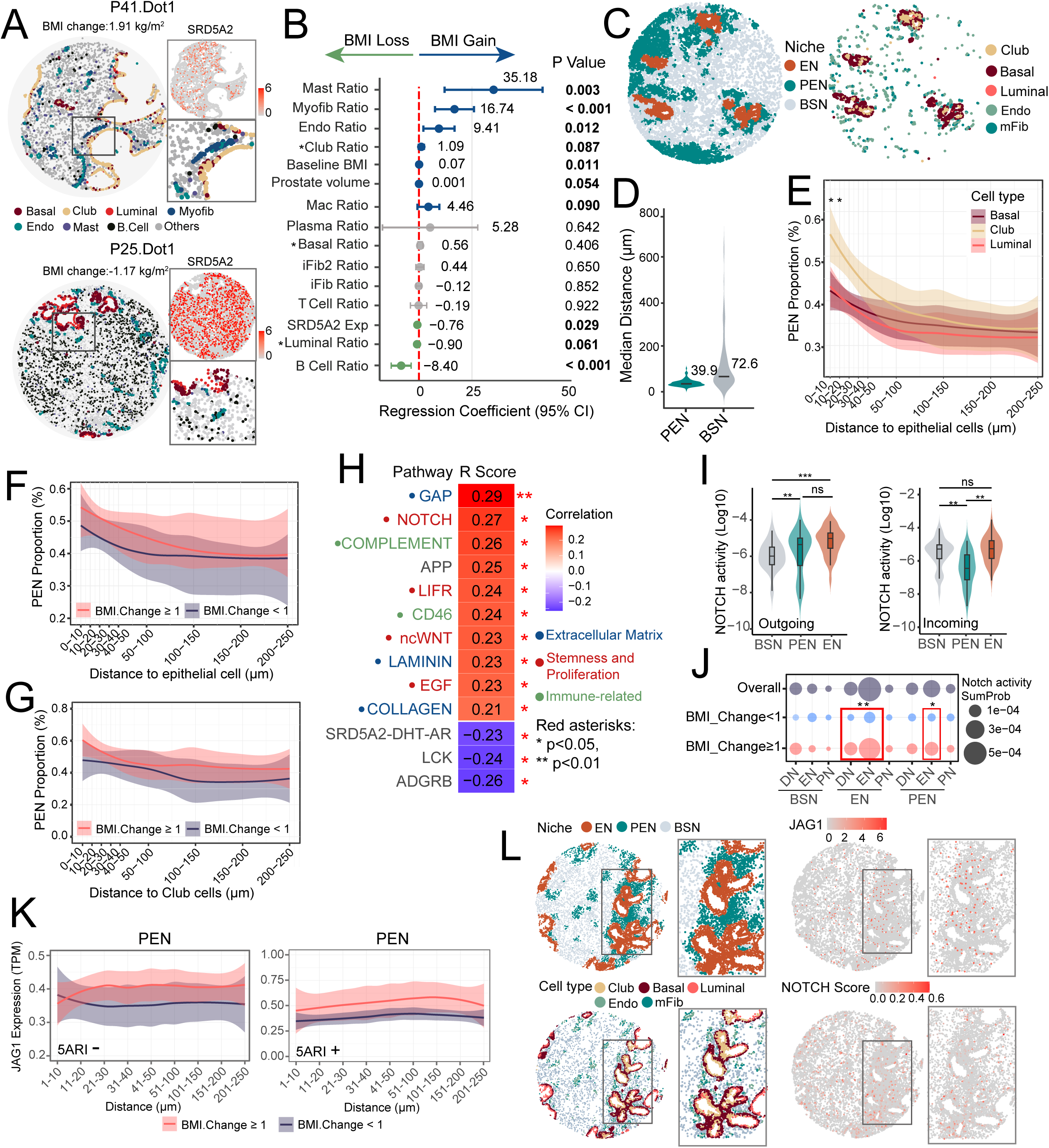
Body weight gain expands stem-like epithelium via NOTCH-enriched niches. The cell clustering and annotation of Xenium data, are provided in Supplementary Methods and Figure S6A–B. **A**: Representative regions showing cellular composition, spatial localization, and SRD5A2 expression across two tissues. **B**: Forest plot depicting factors associated with BMI changes in linear regression analysis. * For linear regression analysis between epithelial cells and BMI change, regions with epithelial cell proportion >5% were selected, resulting in a total of 88 regions included in the analysis. **C**: Left: Representative regions showing the spatial distribution of the three functional niches (EN: epithelial niche; PEN: peri-epithelial niche; BSN: background stromal niche). Right: Corresponding distributions of epithelial (Club, Basal, Luminal), Endo, and mFib cells in the representative regions. **D**: Violin plot showing differences in the median distance of the PEN and BSN niches to epithelial cells. Lines indicate the median values for each group. **E**: LOESS-smoothed curve displays the average proportions of PEN across varying spatial distances from three epithelial cell types, Club, Basal, and Luminal. *P < 0.05. **F-G**: LOESS-smoothed PEN proportions across spatial distances from epithelial (F) and club cells (G) in BMI change ≥1 vs. <1 groups in patients without 5ARI. **H**: CellChat calculated pathway strengths for each region; heatmap shows correlation coefficients (R scores) of pathways significantly associated with BMI change. *P < 0.05, ** P < 0.01. **I**: Violin plots showing NOTCH pathway outgoing (left) and incoming (right) strengths for different functional niches. **J**: Dot plot showing NOTCH pathway interaction strengths between functional niches across BMI change groups and overall. *P < 0.05, ** P < 0.01. **K**: LOESS-smoothed expression of the NOTCH ligand JAG1 across spatial distances from epithelial cells BMI change ≥1 vs. <1 groups, stratified by 5ARI treatment (Left: 5ARI-; Right: 5ARI+). TPM: Transcripts Per Million. **L**: Representative regions of sample with BMI gain (2.03 Kg/m^2^) showing spatial distributions of niches, cell types, JAG1 expression, and NOTCH pathway scores across functional niches. The PEN niche was predominantly located in epithelium with high club cell abundance and exhibited elevated JAG1 expression and NOTCH activity. iFib: Inflammatory Fibroblast; Endo: Endothelial cell; Myofib: Myofibroblast; Endo: Endothelial cell; Mac: Macrophage.

To define spatially integrated cellular and molecular units, we classified BPH tissues into epithelial niche (EN), peri-epithelial niche (PEN) and background stromal niche (BSN) using established spatial and molecular criteria [23] (Figure 5C). EN contained predominantly epithelial cells, whereas PEN and BSN were stromal-derived (Figure 5C, Suppl Figure 7A). PEN was enriched for growth/stemness pathways (e.g., NOTCH) and localized adjacent to the epithelium with higher Endo and mFib content, while BSN showed extracellular matrix signatures and diffuse distribution (Figure 5C-D, Suppl Figure 7A-B). Notably, PEN preferentially surrounded stem-like club cells over luminal or basal cells (Figure 5E; Suppl Figure 7D). Patients with BMI gain (≥1 kg/m²) exhibited increased periepithelial and periclub cell PEN abundance, independent of 5ARI treatment (Figure 5F-G, Suppl Figure 7E-F).

To delineate PEN’s role in BMI-associated epithelial stemness, we analyzed cell-cell signaling. BMI gain correlated positively with growth/stemness pathways (e.g., NOTCH) and inversely with SRD5A2-DHT-AR axis (Figure 5H), with NOTCH activation unaffected by 5ARI treatment (Suppl Figure 7G). PEN and EN emerged as major NOTCH signal sources, with EN also the principal recipient (Figure 5I). At the cellular level, club cells, mFib, Endo, and iFib acted as key Notch sources, while club cells as the main recipients (Suppl Figure 7H). EN-to-EN and PEN-to-EN interactions, particularly JAG1-NOTCH1/2, were significantly enhanced with BMI gain (Figure 5J; Suppl Figure 7I). Spatial maping revealed BMI gain increased JAG1 expression in PEN adjacent to epithelium, suggesting establishment of a PEN-based NOTCH niche that promotes epithelial stemness (Figure 5K, L).

### Body weight control improves the therapeutic efficacy of 5ARI

Collectively, these results indicate that prostate enlargement induced by HFD or elevated BMI occurs independently of SRD5A2 signaling - the primary pathway targeted by 5ARI treatment. This prompted us to investigate whether prostate enlargement driven by body weight gain responds to 5ARI therapy. Multiple restricted cubic spline analysis showed that increases in BMI change were associated with greater prostate volume growth rates in both 5ARI-treated and untreated patients. In the 5ARI-treated patients, 5ARI was effective in reducing prostate volume when patient BMI change was low or negative, but the treatment efficacy diminished with BMI change. (Figure 6A). Subsequently, we performed stratified analyses on the cohort to assess potential efficacy variations across different BMI change categories. Patients were classified into High BMI increase versus Low BMI increase categories using multiple cutoff values. Treatment efficacy was operationally defined as achieving > 35% reduction in prostate volume growth rate. Figure 6C showed that as the cutoff value increased, the treatment efficacy of 5ARI declined in the High-BMI-increase group but remained stable in the Low-BMI-increase group (Figure 6B). The treatment efficacy became non-significant in the High-BMI-increase group at the cutoff value of 2.48 (95% CI included 0). Subsequent stratification at this threshold demonstrated 59% ([OR_low_-OR_high_]/OR_low_) relative efficacy reduction in the High-BMI-increase group versus Low-BMI-increase group (Figure 6C). Moreover, we further investigated whether substantial weight gain affects the efficacy of 5ARIs on AUA Symptom Score (AUASS). Multivariable analysis revealed although 5ARIs were linked to greater AUASS reduction (Suppl Figure 8A), the treatment efficacy became insignificant in the High-BMI-increase group (Suppl Figure 8B). These findings suggest that when weight gain becomes particularly substantial, weight gain-induced SRD5A2-independent growth pathways may dominate over the SRD5A2-dependent mechanisms targeted by 5ARIs.

**Figure 6:**
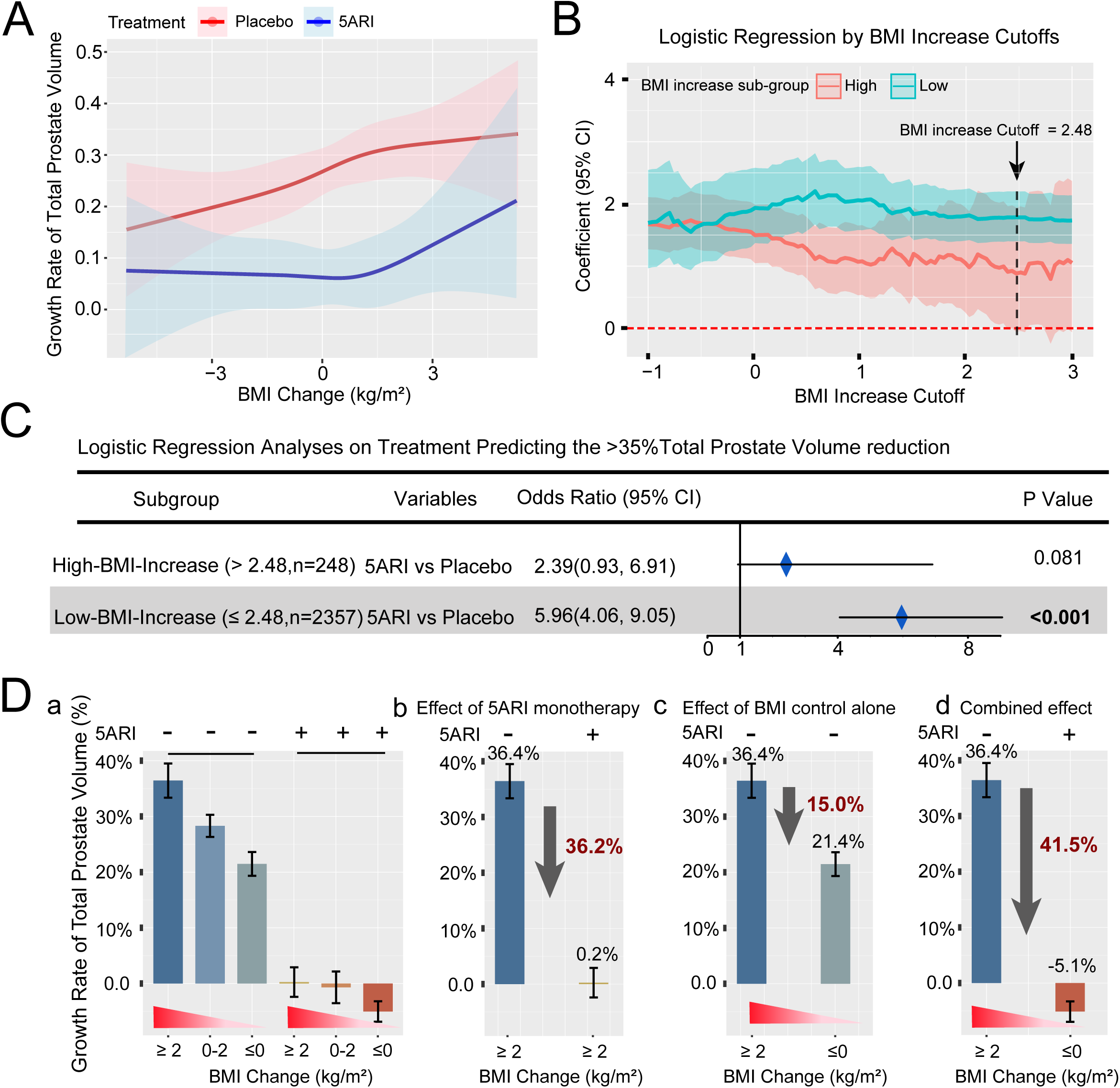
The effect of body weight control on the therapeutic efficacy of 5ARI. **A**: Multiple restricted cubic spline analysis showing the association between BMI change and growth rate of total prostate volume in the 5ARI-treated and placebo groups. The included variables were age, AUA Symptom Score, BMI, transition zone index, and α-blocker status. **B**: Across a range of cut-off values, patients were divided into High-BMI-increase and Low-BMI-increase groups. Logistic regression assessed the association between 5ARI therapy and the likelihood of >35% reduction in prostate volume growth. The x-axis shows a range of cutoff values of BMI increase; The y-axis shows the estimated regression coefficients (with 95% confidence intervals) for therapy efficacy in the High-BMI-increase and Low-BMI-increase subgroups at each cutoff point on the x-axis. **C**: Logistic regression analysis of 5ARI therapy efficacy in predicting >35% total prostate volume reduction: Stratified by BMI-increase subgroups (Cutoff = 2.48). **D**: Bar graphs illustrate the prostate volume growth rate across BMI change subgroups (≥2, 0–2, ≤0 kg/m²) with or without 5ARI treatment in the MTOPS cohort (a). Effect of 5ARI monotherapy (b), BMI control alone (c), and combined intervention (d) on prostate volume growth rate.

We then further stratified patients by baseline BMI and subsequent BMI change, finding that 5ARI efficacy was lowest and not significant in those who transitioned from normal to obese (Cohort 2), followed by those with baseline obesity and further weight gain (Cohort 4). Patients who maintained a normal BMI (Cohort 1) showed the greatest benefit (Suppl Figure 9). Finally, we investigated whether weight management affected prostate growth rate and efficacy of 5ARI in the MTOPS cohort (Figure 6D). Patients who exhibited body weight gain (BMI change >2) had a prostate growth rate of 36.4%, compared to 21.4% in those who maintained their weight over time (BMI change ≤0), representing a 15% reduction in prostate growth associated with weight control alone. Among patients receiving 5ARI therapy, prostate growth rate was 0.2%, representing a 36% reduction compared to those not receiving 5ARIs (36.4%). Most notably, patients who both received 5ARI therapy and maintained stable weight (BMI change ≤0) exhibited a growth rate of -5.1%, amounting to a 41% reduction compared to patients with weight gain and no 5ARI use. These findings suggest that weight control combined with 5ARI use had the maximum benefit in reducing prostate growth rate.

## Discussion

The prostate exhibits pronounced spatial heterogeneity. The proximal region, which corresponds to transition/central zone, area responsible for bladder outlet obstruction leading to lower urinary tract symptoms associated with BPH [24], is notably enriched for stem-like, club-like epithelial cells that mirror pulmonary club cells in both morphology and transcriptomic profiles [24,25]. These proximal urethral club-like cells display reduced AR signaling and increased expression of luminal progenitor markers [24–29], show enhanced survival in 3D spheroid culture, and retain regenerative capacity across castration-regeneration cycles [28]. Moreover, club-like cells within atrophic prostate regions drive tissue regeneration following withdrawal of 5ARIs [29].

Building on these insights, our analysis reveals that HFD-induced obesity drives distinct metabolic reprogramming in the proximal urethral epithelium. This metabolic stress is marked by an expansion of stem-like urethral club cells, accompanied by suppressed expression of SRD5A2 expression and attenuated AR signaling. Notably, this HFD-driven expansion of progenitor-like cells persists in *Srd5a2*-null mice and in BPH patients treated with 5ARIs, underscoring a bypass mechanism that operates independently of canonical SRD5A2-dependent androgen signaling [16; 30]. This androgen-to-stemness transition is reminiscent of changes observed under therapeutic androgen blockade (such as 5ARI administration or androgen deprivation therapy), wherein luminal prostate epithelial cells adopt a club cell-like progenitor state when AR signaling is inhibited [28,30]. In our study, obesity-related metabolic stress elicited a similar form of epithelial plasticity, suggesting a broader adaptive program in which prostatic epithelial proliferation shifts from androgen dependence to stemness-driven growth under stress conditions [25]. These insights may help explain mechanisms of resistance to androgen deprivation in advanced prostate cancer, highlighting that a patient’s metabolic status can significantly influence therapeutic response [31, 15].

In addition to intrinsic reprogramming, obesity reorganizes the spatial distribution of the stromal niche and rewires stromal-epithelial crosstalk. Our analyses reveal that club-like cells receive disproportionately strong stromal-derived inputs relative to other epithelial populations, which are most prominently NOTCH signaling and reflecting their reliance on niche cues than on steroid hormones. This dependence is further amplified by obesity. Given the established roles of Notch in epithelial stemness [32,33], and in prostatic overgrowth and resistance to 5ARIs [34], obesity may promote the expansion of proximal club-like cells by reinforcing NOTCH-mediated stromal-epithelial interactions within the niche.

Prior clinical data indicate that obesity accelerates prostate growth and blunts the prostate volume (PV) reduction achievable with 5ARI [35]. In our and MTOPS cohort, however, longitudinal BMI gain, rather than baseline BMI, emerged as the dominant anthropometric driver of increases in total prostate volume and transition zone volume These observations align with life⍰course data showing that becoming overweight/obese (i.e., category transitions), rather than small weight changes, elevates the risk of clinically enlarged prostates (≥30 cc) and with the central role of transition zone expansion in age⍰related prostatic enlargement [36].

Mechanistically, dynamic weight gain plausibly activates multiple signaling pathways, including the NOTCH pathway, promote prostate growth by triggering an androgen-to-stemness transition growth program, whereas stable adiposity may reflect a metabolically adapted state with less trophic signaling flux.

Stratification by BMI change further revealed diminished responsiveness to 5ARI therapy among men with the greatest BMI gains mirroring prior reports that obesity attenuates prostate volume reduction with 5ARIs [37,38]. This blunting of 5ARI efficacy is biologically coherent with obesity⍰associated hypermethylation of the SRD5A2 promoter and reduced SRD5A2 protein, together with a shift toward SRD5A2⍰independent, stemness⍰driven growth programs observed in diet⍰induced prostate inflammation [39,40]. Collectively, our findings highlight the limitations of 5ARI monotherapy in the setting of metabolic dysfunction and support a more personalized approach that considers weight trajectory and molecular context (e.g., SRD5A2 status).

## Conclusion

Our findings demonstrate that obesity promotes proximal prostate enlargement through an SRD5A2-independent, stem cell-driven mechanism that is unresponsive to 5ARI therapy alone. In obese patients, combining pharmacological treatment with weight management yields synergistic effects on reducing prostate volume growth, highlighting the need for metabolically informed, personalized approaches to BPH management.

## Acknowledgements

We gratefully acknowledge Dr. Chad Vezina’s laboratory at the University of Wisconsin– Madison for generously providing the *Srd5a2*^+/–^ mice used for breeding. We thank the Tissue Microarray Lab & Single Cell and Spatial Lab at Washington University School of Medicine for assistance in processing tissue samples for TMA and Xenium spatial transcriptomic sequencing. We also thank Dr. Linus Tsai at Beth Israel Deaconess Medical Center for performing the single-cell RNA sequencing at the Boston Nutrition and Obesity Center Functional Genomics and Bioinformatics Core. The MTOPS study was conducted by its investigators and supported by the National Institute of Diabetes and Digestive and Kidney Diseases (NIDDK). Data and biospecimens from the MTOPS study used in this work were obtained from the NIDDK Central Repository. This manuscript was not prepared in collaboration with the MTOPS investigators and does not necessarily reflect the views or opinions of the MTOPS study, the NIDDK Central Repository, or the NIDDK.

## Authors’ contributions

ZW and AO performed study design, conceptualization, and supervised this research. ZW, XL and CS designed, executed, and interpreted the experiments on mouse models. RG reviewed the slides for TMA. XL and SW conducted scRNA-seq and Xenium spatial transcriptomic sequencing data analysis. XL conducted MTOPS clinical data and sequencing data analysis. AF reviewed clinical study protocols and edited the manuscript. RK, BG and HR collected the biorepository specimens and edited the manuscript. CS and YT conducted and interpreted animal study and IHC experiments. XL, CS, ZW and BG analyzed data, designed figures, and wrote the manuscript. ZW and AO reviewed and edited the manuscript. All authors reviewed and approved the manuscript.

## Ethics Approval and Consent to Participate

Research involving human tissues from the MTOPS study and our institutional cohort was conducted using de-identified patient samples. Our study using biorepository was approved by the Institutional Review Board at Beth Israel Deaconess Medical Center (IRB#: 2019P000728). All animal experiments were conducted in accordance with the U.S. NIH Guide for the Care and Use of Laboratory Animals and were approved by IACUC at BIDMC.

## Data Availability Statement

The scRNA-seq data are available in NCBI’s GEO (https://www.ncbi.nlm.nih.gov/geo) under accession numbers GSM7504456 (M1), GSM7504457 (M2), GSM8117423 (M3), GSM8117425 (M4), GSM7504455 (M5), GSM8117424 (M6), GSM8117426 (M7). Raw data of quantification in figures is available in the supplemental Supporting Data Values file. Paired RNA-Seq data from 108 baseline/follow-up biopsies from the MTOPS cohort were downloaded from Gene Expression Omnibus (GEO) (GSE262946) [18]. Xenium spatial transcriptomics data, further information, results, reagents, and other supporting data used or analyzed in this study are available from the corresponding author upon reasonable request.

## Reference

[1] Okunogbe, A., Nugent, R., Spencer, G., Powis, J., Ralston, J. & Wilding, J. Economic impacts of overweight and obesity: current and future estimates for 161 countries. BMJ Glob. Health 7, e007163 (2022).

[2] GBD 2019 Risk Factors Collaborators. Global burden of 87 risk factors in 204 countries and territories, 1990–2019: a systematic analysis for the Global Burden of Disease Study 2019. Lancet 396, 1223–1249 (2020).

[3] Gacci, M., Corona, G., Vignozzi, L. et al. Metabolic syndrome and benign prostatic enlargement: a systematic review and meta-analysis. BJU Int. 115, 24–31 (2015).

[4] Chen, X., Yang, S., He, Z. et al. Comprehensive analysis of the global, regional, and national burden of benign prostatic hyperplasia from 1990 to 2021. Sci. Rep. 15, 5644 (2025).

[5] Chang, K., Li, B., Wang, G., Zhou, H., Chen, Y. & Gu, H. Association between Chinese visceral adiposity index and lower urinary tract symptoms suggestive of benign prostatic hyperplasia (LUTS/BPH): a national cohort study. BMC Urol. 25, 15 (2025).

[6] Kristal, A. R., Arnold, K. B., Schenk, J. M. et al. Race/ethnicity, obesity, health-related behaviors and the risk of symptomatic benign prostatic hyperplasia: results from the Prostate Cancer Prevention Trial. J. Urol. 177, 1395–1400 (2007).

[7] Parsons, J. K., Messer, K., White, M. et al. Obesity increases and physical activity decreases lower urinary tract symptom risk in older men: the Osteoporotic Fractures in Men study. Eur. Urol. 60, 1173–1180 (2011).

[8] Vignozzi, L., Gacci, M. & Maggi, M. Lower urinary tract symptoms, benign prostatic hyperplasia and metabolic syndrome. Nat. Rev. Urol. 13, 108–119 (2016).

[9] Penson, D. F., Munro, H. M., Signorello, L. B. et al. Obesity, physical activity and lower urinary tract symptoms: results from the Southern Community Cohort Study. J. Urol. 186, 2316–2322 (2011).

[10] Fu, X., Wang, Y., Lu, Y., Liu, J. & Li, H. Association between metabolic syndrome and benign prostatic hyperplasia: the underlying molecular connection. Life Sci. 358, 123192 (2024).

[11] Negi, S. K., Desai, S., Faujdar, G. et al. The correlation between obesity and prostate volume in patients with benign prostatic hyperplasia: a prospective cohort study. Urologia 91, 512–517 (2024).

[12] Zaza, M. M., Salem, T. A., Hassanin, I. S. F. & Soliman, M. H. A. Effect of body mass index on prostate volume and prostate-specific antigen in men over 50: a cross-sectional study. Urologia 90, 224–229 (2023).

[13] Ge, R., Wang, Z., Bechis, S. K. et al. DNA methyltransferase 1 reduces expression of SRD5A2 in the aging adult prostate. Am. J. Pathol. 185, 870–882 (2015).

[14] Xue, B., Wu, S., Sharkey, C. et al. Obesity-associated inflammation induces androgenic to estrogenic switch in the prostate gland. Prostate Cancer Prostatic Dis. 23, 465–474 (2020).

[15] Kwon, O. J., Zhang, B., Zhang, L. & Xin, L. High fat diet promotes prostatic basal-to-luminal differentiation and accelerates initiation of prostate epithelial hyperplasia originated from basal cells. Stem Cell Res. 16, 682–691 (2016).

[16] Sharkey, C., Long, X., Al-Faouri, R. et al. Enhanced prostatic Esr1(+) luminal epithelial cells in the absence of SRD5A2. J. Pathol. 263, 300–314 (2024).

[17] McConnell, J. D., Roehrborn, C. G., Bautista, O. M. et al. The long-term effect of doxazosin, finasteride, and combination therapy on the clinical progression of benign prostatic hyperplasia. N. Engl. J. Med. 349, 2387–2398 (2003).

[18] Choi, H. Y., Torkko, K. C., Lucia, M. S. et al. Change in prostate tissue gene expression following finasteride or doxazosin administration in the medical therapy for prostatic symptoms (MTOPS) study. Sci. Rep. 14, 19164 (2024).

[19] Sharkey, C., Long, X., Wang, Z. et al. Zonal growth pattern of the prostate is affected by age and body mass index. J. Urol. 207, 876–884 (2022).

[20] Joseph, D. B., Henry, G. H., Malewska, A. et al. Urethral luminal epithelia are castration-insensitive cells of the proximal prostate. Prostate 80, 872–884 (2020).

[21] Vellky, J. E., Wu, Y., Moline, D. et al. Single-cell RNA sequencing of human prostate basal epithelial cells reveals zone-specific cellular populations and gene expression signatures. J. Pathol. 262, 212–225 (2024).

[22] Chen, W., Pascal, L. E., Wang, K. et al. Differential impact of paired patient-derived BPH and normal adjacent stromal cells on benign prostatic epithelial cell growth in 3D culture. Prostate 80, 1177–1187 (2020).

[23] Vannan, A., Lyu, R., Williams, A. L. et al. Spatial transcriptomics identifies molecular niche dysregulation associated with distal lung remodeling in pulmonary fibrosis. Nat. Genet. 57, 647–658 (2025).

[24] Henry, G. H., Malewska, A., Joseph, D. B. et al. A cellular anatomy of the normal adult human prostate and prostatic urethra. Cell Rep. 25, 3530–3542.e3535 (2018).

[25] Huang, F. W., Song, H., Weinstein, H. N. et al. Club-like cells in proliferative inflammatory atrophy of the prostate. J. Pathol. 261, 85–95 (2023).

[26] Fei, X., Liu, J., Xu, J. et al. Integrating spatial transcriptomics and single-cell RNA-sequencing reveals the alterations in epithelial cells during nodular formation in benign prostatic hyperplasia. J. Transl. Med. 22, 380 (2024).

[27] Kwon, O. J., Choi, J. M., Zhang, L. et al. The Sca-1(+) and Sca-1(–) mouse prostatic luminal cell lineages are independently sustained. Stem Cells 38, 1479–1491 (2020).

[28] Karthaus, W. R., Hofree, M., Choi, D. et al. Regenerative potential of prostate luminal cells revealed by single-cell analysis. Science 368, 497–505 (2020).

[29] Huang, F. W., Song, H., Weinstein, H. N. et al. Club-like cells in proliferative inflammatory atrophy of the prostate. J. Pathol. 261, 85–95 (2023).

[30] Joseph, D. B., Henry, G. H., Malewska, A. et al. 5-Alpha reductase inhibitors induce a prostate luminal-to-club cell transition in human benign prostatic hyperplasia. J. Pathol. 256, 427–441 (2022).

[31] MustaLă, T., Jinga, D. C., Lazăr, I., Martin, S. C., Sîrbu, A. E. & Fica, S. The impact of obesity on prostate cancer and progression to castration resistance: real-world data from a Romanian center. J. Clin. Med. 14, 3146 (2025).

[32] Bigas, A. & Porcheri, C. Notch and stem cells. Adv. Exp. Med. Biol. 1066, 235–263 (2018).

[33] Centonze, A., Lin, S., Tika, E. et al. Heterotypic cell-cell communication regulates glandular stem cell multipotency. Nature 584, 608–613 (2020).

[34] Jin, R., Forbes, C. M., Miller, N. L. et al. Transcriptomic analysis of benign prostatic hyperplasia identifies critical pathways in prostatic overgrowth and 5-alpha reductase inhibitor resistance. Prostate 84, 441–459 (2024).

[35] Muller, R. L., Gerber, L., Moreira, D. M. et al. Obesity is associated with increased prostate growth and attenuated prostate volume reduction by dutasteride: a randomized controlled trial. Eur. Urol. 63, 1115–1121 (2013).

[36] Khan, S., Wolin, K. Y., Pakpahan, R. et al. Body size throughout the life-course and incident benign prostatic hyperplasia-related outcomes and nocturia. BMC Urol. 21, 47 (2021).

[37] Muller, R. L., Gerber, L., Moreira, D. M. et al. Obesity is associated with increased prostate growth and attenuated prostate volume reduction by dutasteride: a randomized controlled trial. Eur. Urol. 63, 1115–1121 (2013).

[38] Parsons, J. K., Schenk, J. M., Arnold, K. B. et al. Finasteride reduces the risk of incident clinical benign prostatic hyperplasia: a randomized controlled trial. Eur. Urol. 62, 234– 241 (2012).

[39] Bechis, S. K., Otsetov, A. G., Ge, R. et al. Age and obesity promote methylation and suppression of 5α-reductase 2: implications for personalized therapy of benign prostatic hyperplasia. J. Urol. 194, 1031–1037 (2015).

[40] Lin, Z.-M., Fan, D.-D., Jin, S., Liu, Z.-L. & Niu, Y.-N. Methylated CpG dinucleotides in the 5-α reductase 2 gene may explain finasteride resistance in benign prostatic enlargement patients. Asian J. Androl. 23, 266–272 (2021).

