## Supplemental Methods for "Stem-like Prostate Remodeling in Obesity Mediates Resistance to 5α-Reductase Inhibition Therapy in BPH"

**Supplementary Methods**

**Mouse model generation**

All animal procedures were approved by the Beth Israel Deaconess Medical Center IACUC (protocol number: 026-2019). The detailed generation process has been described in our previous publication[1]. Briefly Srd5a2-null mice were generated using *Srd5a2*^creErt2^ knock-in alleles (*Srd5a2*^G2aCE^ knock-in allele; Jackson Laboratory, Bar Harbor, ME, USA), which disrupt endogenous Srd5a2 while expressing EGFP and Cre-ERT2 in the male reproductive tract. Heterozygous (*Srd5a2*^+/−^) males were bred to WT C57BL/6 females, and subsequent intercrosses produced homozygous knockouts (*Srd5a2*^−/−^). WT and heterozygous littermates showed no phenotypic differences and served as controls.

**Dietary intervention**

*Srd5a2*^+/−^ and *Srd5a2*^−/−^ mice were fed with either a regular-fat diet (RFD) (RMH3000:14% fat, 60% carbohydrates, 26% protein; Lab Diet, St Louis, MO) or a high-fat diet (D12451: 45% fat, 35% carbohydrates, 20% protein; Research Diets, New Brunswick, NJ) for 3 months.

**Clinical parameter definition**: BMI change was calculated as change in BMI from baseline to last follow-up; Prostate volume growth rate was calculated as the change in prostate volume from baseline to last follow-up, divided by the baseline prostate volume.

**Clinical data analyses**

Categorical data were expressed as count and percentage and continuous data were expressed as mean ± standard deviation (SD). The *χ*^2^-test or Fisher’s exact test was used for categorical data when appropriate. Depending on the data distribution, either the Student’s t-test or the Wilcoxon test was used for continuous variables. Linear and logistic regression were performed for univariate and multivariable linear model analyses. In the linear regression analyses, the regression coefficients were uniformly multiplied by 100 in Figures 4, 6, and Suppl Figure 5 to facilitate visualization, as their original values were relatively small. Restricted cubic spline analysis with four knots was employed to model potential non-linear relationships. LOESS curve was used for nonlinear fitting of two continuous variables. Pearson’s correlation was used to evaluate the correlation between two continuous variables. For all statistical analyses, *P* < 0.05 was considered statistically significant. Statistical tests for every figure are justified as appropriate and all data meet the assumptions of the responding tests.

**Prostate isolation and enzymatic digestion**

Prostates were isolated as previously described [2]. Briefly, dissected prostates were digested in Hank's Balanced Salt Solution (HBSS; Gibco, cat# 24020-117) containing 2 mg/mL collagenase type II (Life Technologies, cat# 17101-015), 10 μM ROCK inhibitor Y-27632 (Abcam, cat# ab120129), 1 nM dihydrotestosterone (Cerilliant, cat# D-073), and 2 U/mL DNase I (Invitrogen, cat# 18068-015) for 2 hours at 37°C. The tissue was then subjected to a second digestion in TrypLE (Life Technologies, cat# 12605-010) supplemented with 10 μM Y-27632 at 37°C for 10 minutes to obtain a single-cell suspension. Red blood cells were lysed using RBC lysis buffer (BioLegend, cat# 420301) for 3 minutes on ice. The cells were then washed twice with 0.02% BSA in PBS and assessed for viability using trypan blue exclusion (Gibco, cat# 15250061). All experiments were conducted using cell preparations with ≥80% viability.

**Single-cell RNA sequencing (scRNA-seq)**

ScRNA-seq was performed on seven fresh prostate tissues from *Srd5a2*^−/−^ and *Srd5a2*^+/−^ mice fed either a RFD or HFD, using the 10x Genomics platform. Libraries were pooled and sequenced on Illumina NextSeq 500 and NovaSeq 6000 platforms. Grouping information and mapping statistics are provided in Supplementary Table S4. Sequencing yielded 50605 cells with a median of 3783 UMI and an averages of 2066 genes per cell.

**Single-cell sequencing and data analysis**

***Mapping of single-cell RNA sequencing data***

The raw reads were mapped to the mouse reference genome (mm10) with Ensembl gene annotation version 92 using CellRanger v3.1.0 (10× Genomics)

***Quality Control***

To ensure high-quality single-cell transcriptomic data, we performed quality control filtering before downstream analysis. Cells with fewer than 500 or more than 7,000 detected genes, or with >10% of reads mapping to mitochondrial genes, were excluded. Genes expressed in fewer than 3 cells were filtered out, and cells with fewer than 500 detected genes were excluded.

***Integrated and Clustering analyses***

We used Seurat (v 5.2.1)[3] to create data objects from the matrix outputs. Data were normalized to log scale using the ‘NormalizeData’ function with a default scale parameter of 10000. The top 2000 highly variable genes were identified using the ‘FindVariableFeatures’ function with the “vst” method with default parameters. Mitochondrial and ribosomal genes were removed if they were highly variable. Cells from the seven samples were then integrated using harmony algorithm[4].

The effects of variation in sequencing depth were regressed out by including ‘nUMI’ as a parameter in the ‘ScaleData’ function. Scaled highly variable genes were used as input for PCA using the ‘RunPCA’ function. The first 15 principal components (PCs) and a resolution of 0.1 were used for clustering using ‘FindClusters’. UMAP was applied for two-dimensional representation of the first 15 PCs with ‘RunUMAP’.

***Epithelial cell type identification***

The SingleR [5] tool (version 2.4.0) was used to annotate epithelial cell types. Our dataset revealed 13 epithelial clusters. To further identify the anatomical subregions of the prostate epithelium, we used the epithelial cell-type expression matrix from the dataset published by Joseph et al. [6] as a reference.

***DEG calculation***

Differentially expressed genes (DEGs) across all clusters and between the HFD and RFD groups were identified using the “FindAllMarkers” function in Seurat, applying the “roc” method for statistical testing. DEGs were defined based on the following criteria: (1) expression in at least 30% of cells within a given cluster; and (2) a p-value < 0.05.

***Pathway or gene set enrichment analysis***

An expression matrix obtained from the data slot of the Seurat object was used for enrichment analysis. Gene Set Variation Analysis (GSVA) [7] was performed to assess pathway activity using publicly available prostate gene sets. The H hallmark gene sets and C2 CP:KEGG gene sets were downloaded from MSigDB v6.2 ([https://www.gsea-msigdb.org/gsea/msigdb/mouse/collections.jsp](https://www.gsea-msigdb.org/gsea/msigdb/mouse/collections.jsp" \t "_new)). Prostate-specific gene signatures and their corresponding literature sources are summarized in Supplementary Table 3. Only genes upregulated in specific cell subtypes were included in the enrichment analysis. For GSVA, statistically significant differences in enrichment scores between groups were determined using the R package limma [8], with a false discovery rate (FDR)-corrected p-value threshold of < 0.05.

***Interaction between cell types***

The expression count matrix of each cell lineage, along with corresponding cell type metadata, was extracted from the Seurat object and used as input for CellPhoneDB (v5)[9], utilizing the ‘cpdb_statistical_analysis_method’ in Python (v3.9.21). CellPhoneDB can only process human genes; therefore, before analysis, the R package homologene was employed to perform homologous gene conversion between mouse and human genes. This analysis was conducted to identify statistically significant receptor–ligand interactions (p < 0.05) by comparing observed expression patterns against a null distribution, based on a curated receptor–ligand interaction database.

***Trajectory analysis***

Single-cell gene expression trajectories were constructed using Monocle (v2.30.0) [10] by importing Seurat data via the ‘importCDS’ function. Size factors and dispersion values were estimated using the ‘estimateSizeFactors’ and ‘estimateDispersions’ functions, respectively. Highly variable genes were identified using the ‘dispersionTable’ function and filtered based on a mean expression threshold of 0.1. These genes were then used as input for the ‘setOrderingFilter’ function. Dimensionality reduction was performed using the ‘reduceDimension’ function with the DDRTree method, and cells were ordered along a pseudotime trajectory using the ‘orderCells’ function.

**Immunostaining**

Immunostaining of mouse prostate tissues, MTOPS biopsy samples, and surgical specimens from the BIDMC biorepository was performed as previously described [1]. Briefly, formalin-fixed, paraffin-embedded tissue sections were subjected to deparaffinization, antigen retrieval, and blocking, followed by incubation with primary antibodies against Trop2 (R&D Systems, AF1122-SP, dilution 1:100), Krt5 (LSBio, C759620, dilution 1:100), SCGB1A1 (Novus Biologicals, MAB4218-SP, dilution 1:100), and SRD5A2 (Invitrogen, PA5-42550, dilution 1:200). Sections were then counterstained to visualize target proteins. Images were acquired from stained sections, and the average immunoreactive score (IRS) was calculated in accordance with the protocol described in the previous study [1].

**Tissue microarray (TMA) construction for Xenium analysis**

To maximize efficiency per run, multiple samples could be placed on a single Xenium slide (10.45 mm × 22.45 mm) using a TMA design. A 5-μm section of each prostate FFPE block was hematoxylin and eosin (H&E) stained and presented to a physician who identified areas of interest and labeled them as stroma or epithelia relative to the sample. Blocks were designed in a 6 × 14 pattern for 1 mm cores, respectively. Sample cores were punched and placed manually using a Beecher Manual Tissue Microarrayer. Empty core spaces were filled with core punches taken from blank paraffin blocks. Once complete, blocks are placed face down on a clean glass slide and briefly heated in a warm drawer (~45 °C) to slightly melt the paraffin together and even the block face. TMA blocks were then cooled to room temperature, removed from the slide, sealed and stored at 4 °C.

**Xenium In Situ spatial transcriptomic profiling**

Xenium In Situ slides (n = 2, 142 TMA regions) were processed according to the manufacturer’s guidelines for TMA samples (PN-1000465, CG000578 Rev A, 10x Genomics). In brief, 10-mm tissue sections from selected regions were mounted within the defined sample frame (10.45 × 22 mm) on Xenium slides. Cassettes (CG000581 Rev C) were then assembled and subjected to probe hybridization (PN-1000671), ligation, and amplification (CG000760 Rev C). Following autofluorescence quenching and nuclear counterstaining, images were captured and analyzed using the Xenium Analyzer (CG000584 Rev K). Regions of interest were manually delineated from the scanned images, and post-run data for each slide were generated using default settings for downstream analysis.

**Xenium data preprocessing**

*Raw transcriptomic profiling data processing*
Raw transcriptomic profiling from the Xenium 50K panel was processed using Xenium Ranger with default parameters (version 1.7.0, 10x Genomics). The resulting raw count matrix was pre-processed using the Seurat package (version 5.0.0) in R (version 4.3.1). Quality control filtering was performed to retain cells that met the following criteria: nCount_Xenium > 12, nFeature_Xenium > 10, a cumulative proportion of high-quality transcripts matching negative control probes, negative control codewords, or unassigned codewords not exceeding 5%, and a nuclear area between 6 and 80 µm². Spatial visualization of sequencing depth was assessed using the ImageFeaturePlot function. Normalization was performed using the SCTransform function (regularized negative binomial regression) with the Xenium assay specified. Principal component analysis (PCA) was conducted on all detected genes, and the first 30 principal components were used for downstream analysis. Uniform Manifold Approximation and Projection (UMAP) was applied for dimensionality reduction. A k-nearest neighbors graph was constructed using the FindNeighbors function, and cell clustering was performed with the FindClusters. The resulting clusters were visualized on the UMAP embedding using the DimPlot function with cluster labels.

*Cell-type annotation and calculation of SRD5A2 expression in each region*

Through dimensionality reduction and clustering, 13 distinct cell clusters were identified. These were annotated according to marker gene expression and spatial distribution (Figure SA-C). Epithelial clusters within the glandular lumen included basal cells (DST), club cells (CP), and luminal cells (ACPP, NKX3-1). Stromal clusters comprised two inflammatory fibroblast subtypes—iFib (PDGFRA) and iFib2 (FGF2)—as well as myofibroblasts (RGS5), endothelial cells (PECAM1), and diverse immune cells scattered throughout the stromal region, including B cells (MS4A1), macrophages (CD68), mast cells (KIT), T cells (CD4), and plasma cells (MZB1). One cluster, which expressed markers of multiple stromal lineages and was uniformly distributed throughout the stromal region, was designated as background stromal cells. Figure SB shows that SRD5A2 is primarily expressed in iFib, iFib2, and myoFib. We calculated the average expression of SRD5A2 across these three cell types to represent SRD5A2 expression in each region.

*Cell fraction quantification*

Since prostate tissue is primarily composed of epithelial acini and stroma, TMA cores were sampled randomly, resulting in variable epithelial area across different regions. Consequently, calculating cell proportions relative to the total cell count in each core could introduce bias. To address this, proportions of epithelial and stromal cells were calculated separately by normalizing each cell type to the total number of epithelial or stromal cells, respectively. For analyses involving epithelial cells, only regions with an epithelial cell fraction greater than 5% were included, yielding a total of 88 regions.

*Niche Identification Analysis*

Cellular niches were identified using an unsupervised neighborhood-based approach adapted from Vannan et al [11]. For each spatial region, cell-level annotations and two-dimensional coordinates were extracted from Seurat objects. Cell types were one-hot encoded, and for each cell, its 25 nearest spatial neighbors were identified using the FNN R package (get.knn function). The neighborhood composition of each cell was then represented as the sum of one-hot vectors across its nearest neighbors, thereby capturing the local cellular context. To minimize bias arising from differences in global cell-type abundance, the neighborhood composition matrix was z-score scaled. We next applied k-means clustering (k = 3, 20 random starts, seed = 42) to this scaled matrix, assigning each cell to one of eight discrete clusters, which we defined as “niches.”

*Spatial Distance Analysis*

Spatial proximity between cell populations was quantified by calculating nearest-neighbor Euclidean distances based on tissue coordinates from Xenium data. For each target cell, the distance to the closest reference cell was identified using the FNN R package, and the median nearest-neighbor distance was used to represent the spatial relationship between the two populations.

*Analysis of Local Niche Composition*
We quantified the microenvironment surrounding specific target cells by calculating the proportions of neighboring cell types within concentric distance bins (e.g., 0-10 µm, 10-50 µm, up to 200-250 µm). For each sample, cell coordinates and annotations were extracted from Seurat objects, and nearest neighbors were identified using the FNN R package. The relative abundance of each niche type was averaged across all target cells to obtain sample-level estimates of local niche composition.

*Cell–cell communication analysis*

Cell–cell communication analysis was performed on each tissue regions with with epithelial cell proportion >5% using the CellChat R package [12]. Normalized Xenium expression data and cell type annotations were used to create a CellChat object with spatial coordinates. The human ligand-receptor database (CellChatDB.human) was applied, and standard workflows were followed to identify overexpressed genes and ligand-receptor interactions, compute communication probabilities, filter low-confidence interactions, and aggregate the signaling network. Spatial communication probabilities were calculated with population-size correction, a 250 µm interaction range, and a 10 µm contact range.

To quantify NOTCH pathway activity, outgoing and incoming signaling strengths were calculated for each cell type by summing communication probabilities across rows and columns, respectively, and ranking them in descending order. This approach allowed identification of key NOTCH signal-sending and signal-receiving cell populations within the spatial tissue context.

Pathway and gene set enrichment analyses of Xenium data were performed using the same approach as in single-cell sequencing.

Reference

1. Sharkey C, Long X, Al-Faouri R, Strand D, Olumi AF, Wang Z: Enhanced prostatic Esr1(+) luminal epithelial cells in the absence of SRD5A2. J Pathol 2024, 263(3):300-314.
2. Drost J, Karthaus WR, Gao D, et al. Organoid culture systems for prostate epithelial and cancer tissue. Nat Protoc. 2016;11(2):347‐358. 10.1038/nprot.2016.006
3. Butler A, Hoffman P, Smibert P, Papalexi E, Satija R. Integrating single-cell transcriptomic data across different conditions, technologies, and species. Nat Biotechnol. 2018; 36(5): 411-420.
4. Korsunsky I, Millard N, Fan J, et al. Fast, sensitive and accurate integration of single-cell data with Harmony. Nat Methods. 2019 Dec;16(12):1289-1296.
5. Aran D, Looney AP, Liu L, et al. “Reference-based analysis of lung single-cell sequencing reveals a transitional profibrotic macrophage”. Nat Immunol. 2019; 20(2): 163-172.
6. Joseph DB, Henry GH, Malewska AU, et al. rethral luminal epithelia are castration-insensitive cells of the proximal prostate.Prostate. 2020 Aug;80(11):872-884.
7. Hänzelmann S, Castelo R, Guinney J. GSVA: gene set variation analysis for microarray and RNA-seq data. BMC Bioinformatics. 2013; 14: 7.
8. Ritchie ME, Phipson B, Wu D, et al. “limma powers differential expression analyses for RNA-sequencing and microarray studies” . Nucleic Acids Res. 2015; 43(7): e47.
9. Troulé K, Petryszak R, Cakir B et al. CellPhoneDB v5: inferring cell-cell communication from single-cell multiomics data. Nat Protoc. 2025 Mar 25.
10. Patel AP, Tirosh I, Trombetta JJ, et al. Single cell RNA-seq highlights intratumoral heterogeneity in primary glioblastoma. Science. 2014; 344(6190): 1396 -1401.
11. Vannan A, Lyu R, Williams AL, et al. Spatial transcriptomics identifies molecular niche dysregulation associated with distal lung remodeling in pulmonary fibrosis. Nat Genet 2025, 57(3):647-658.
12. Jin S, Plikus MV, Nie Q. CellChat for systematic analysis of cell-cell communication from single-cell transcriptomics. Nat Protoc 2025, 20(1):180-219.
