## Supplemental Figure and Table Legend for "Stem-like Prostate Remodeling in Obesity Mediates Resistance to 5α-Reductase Inhibition Therapy in BPH"

**Supplemental Figure and Table Legends**

**Supplementary Figure 1: A:** Bar graph showing absolute body weights in *Srd5a*2^-/-^ (knockout), *Srd5a2*^+/-^ (heterozygous) and *Srd5a2*^+/+^ (wild-type) mice under regular fat diet (RFD) and HFD conditions. **B**: Comparative schematic of mouse and human prostate anatomy illustrating proximal and distal regions. The left panel shows the murine prostate lobes. analogous proximal (urethra [UR], and ventral prostate [VP]) and distal (anterior prostate [AP], dorsal-lateral prostate [DLP]) regions. The right panel shows a human prostate section with analogous proximal (periurethral and transitional zone) and distal (peripheral zone) regions. Red and green lines indicate proximal and distal regions, respectively.

**Supplementary Figure 2:** **A**: Fishplot showing pseudotime distribution of epithelial cells under HFD vs RFD in *Srd5a2*^-/-^ mice. **B**: Dot plot showing expression of representative AR lineage and Stem genes across proximal epithelial cells in *Srd5a2*^+/-^ mice. **C**: LOESS-fitted average expression of ESR1 target, lineage, and stem gene signatures across pseudotime in estrogen and AR lineage trajectories under *Srd5a*2^-/-^ condition, comparing HFD and RFD groups. **D**: Gene Set Variation Analysis (GSVA)-based comparison of pathway activity scores between HFD and RFD across epithelial cell types in *Srd5a2*^-/-^ mice, shown as t-values from a linear model. **E**: Dot plot showing expression of representative AR lineage and stem genes across proximal epithelial cells in *Srd5a2*^-/-^ mice. **F**: Immunofluorescence staining of Ur.LE marker Trop2 and basal cell marker CK5 in prostates of *Srd5a2*^-/-^ mice under HFD and RFD. **G**: Bar graph showing the ratio of urethral/proximal prostate to total prostate (Trop2^+^ region/whole prostate region) in *Srd5a2*^+/-^ and *Srd5a2*^-/-^ mice under RFD and HFD conditions. **H**: Fold-change in the ratio of urethral/proximal prostate region to total prostate region (HFD vs. RFD) within each genotype.

**Supplementary Figure 3**: **A**: Dot plot illustrating epithelial cell-specific ligand-receptor signaling via AR and ESR1 from stromal cells and lymphocytes in *Srd5a2*^+/−^ and *Srd5a2*^−/−^ mice under RFD and HFD conditions. **B**: Dot plot showing that Srd5a2 is predominantly expressed in prostate fibroblasts cells, and comparing its expression in fibroblasts cells from *Srd5a2*^+/−^ and *Srd5a2*^−/−^ mice under RFD and HFD conditions. **C**: Dot plot showing the Hsd17b12 expression across the cell types and comparing its expression in macrophages and neuron cells from *Srd5a2*^+/−^ and *Srd5a2*^−/−^ mice under RFD and HFD conditions. Fib: Fibroblast; Endo: Endothelial cell; Myofib: Myofibroblast; Mac: Macrophage; NK: Natural killer cell; AP: anterior prostate; DLP: dorsolateral prostate; VP: ventral prostate; Ur: urethral prostate. LE: luminal epithelial cell.

**Supplementary Figure 4**: **A**: Dot plot showing epithelial cell-specific ligand-receptor interactions involving Wnt and Notch pathways from stromal cells in *Srd5a2*^+/−^ and *Srd5a2*^−/−^ mice under RFD and HFD conditions. **B**: LOESS-smoothed Gene Set Variation Analysis (GSVA) scores of Wnt/Notch pathways and receptor expression (TPM: Transcripts Per Million) along pseudotime in *Srd5a2*^−/−^ mice under HFD and RFD.

**Supplementary Figure 5**: **A-C**: Univariate linear regression analyses on factors predicting the growth rate of total prostate volume (A), prostate transition zone volume (B) and prostate peripheral zone volume (C).

**Supplementary Figure 6: A:** Left: UMAP plot presenting the distribution of 13 major cell clusters. Right: Heatmap showing representative marker genes across 13 cell clusters. **B**: Representative regions showing H&E staining (left) and corresponding cell-annotated Xenium spatial transcriptomic images (right). **C**: Linear regression interaction analysis tested whether 5ARI modified the effect of BMI change on other variables. **D**: Immunohistochemistry analysis showing the correlation between BMI change and SRD5A2 expression in BPH tissues from our institution. Left: Representative Immunohistochemistry staining; Right: correlation curve. **E**: Left: LOESS curves showing correlations between BMI change and changes in ssGSEA (Single-sample Gene Set Enrichment Analysis) scores for Club epithelial cells (CE), Hillock epithelial cells (HE), and stem signatures (follow-up values minus baseline values). The ssGSEA scores were derived from RNA-seq data of paired baseline and follow-up prostate biopsy samples from 103 MTOPS cohort patients. Right: Correlation between BMI change and changes in ssGSEA scores stratified by 5ARI treatment. **F**: Left: LOESS curves showing correlations between BMI change and changes in ssGSEA scores for luminal epithelial cell (LE), basal epithelial cell (BE), and AR lineage gene signatures. The ssGSEA scores were derived from RNA-seq data of paired baseline and follow-up prostate biopsy samples from 103 MTOPS cohort patients. Right: Correlation between BMI change and changes in ssGSEA scores stratified by 5ARI treatment. **G**: Correlation between BMI change and DHT change (follow-up values minus baseline values) in MTOPS patients. Colors indicate 5ARI status.

iFib: Inflammatory fibroblast; Endo: Endothelial cell; Myofib: Myofibroblast; Endo: Endothelial cell; Mac: Macrophage.

**Supplementary Figure 7: A:** Bar plot showing cell type proportions across different functional niches. * Significant difference between groups (P < 0.05). **B**: Gene Set Variation Analysis (GSVA) based comparison of pathway activity scores between PEN and BSN. **C**: Dot plot showing expression of representative pathway genes between PEN and BSN. **D**: Violin plot showing median PEN distances to different epithelial cells across regions. **E, F**： LOESS-smoothed PEN proportions across spatial distances from epithelial (E) and club cells (F) in 5ARI-treated patients, stratified by BMI change ≥1 vs. <1. **G**：Linear regression interaction analysis assessed whether 5ARI influenced the effect of BMI change on pathway activity. **H**: Violin plots showing NOTCH pathway outgoing (upper) and incoming (down) strengths across different cell types. **I**: Dot plot showing NOTCH pathway ligand-receptor interaction strengths between functional niches across BMI change groups, with red boxes highlighting pairs exhibiting significant differences.

EN: epithelial niche; PEN: peri-epithelial niche; BN: background stromal niches; iFib: Inflammatory fibroblast; Endo: Endothelial cell; Myofib: Myofibroblast; Endo: Endothelial cell; Mac: Macrophage.

**Supplementary Figure 8: A:** Multivariate Linear Regression Analyses on Factors Predicting the AUA Symptom Score Change. **B**: Multivariate linear regression analyses of factors associated with changes in AUA Symptom Score in the high BMI increase (upper) and low BMI increase (lower) subgroups, respectively.

**Supplementary Figure 9:** Forest plot showing odds ratios (95% CI) from logistic regression analyses assessing the association between BMI status, BMI change, and treatment response to 5α-reductase inhibitors (5ARIs). Patients were stratified by baseline BMI (<24 vs. ≥24) and follow-up BMI or BMI change. Cohort 1: baseline BMI <24 and follow-up BMI ≤24; Cohort 2: baseline BMI <24 and follow-up BMI >24; Cohort 3: baseline BMI ≥24 with BMI change ≤0; Cohort 4: baseline BMI ≥24 with BMI change >0. Odds ratios are plotted on a logarithmic scale.

**Supplementary Table 1:** Clinical demographic information of the MTOPS population included in the study (n = 2605).

**Supplementary Table 2:** Clinical demographic information of the samples used for Xenium spatial transcriptomics.

**Supplementary Table 3:** Gene sets used in the article.

**Supplementary Table 4:** scRNA-seq sample information.
