## Supplementary figures and images for "Stem-like Prostate Remodeling in Obesity Mediates Resistance to 5α-Reductase Inhibition Therapy in BPH"

### Supplemental Figure 1

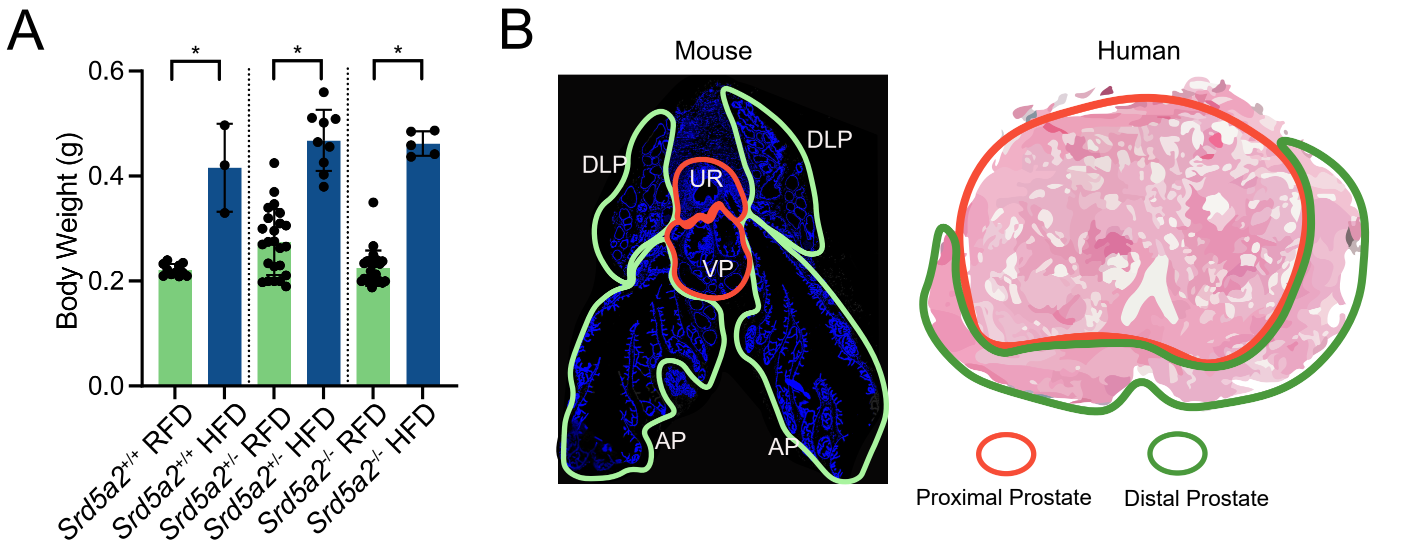

### Supplemental Figure 2

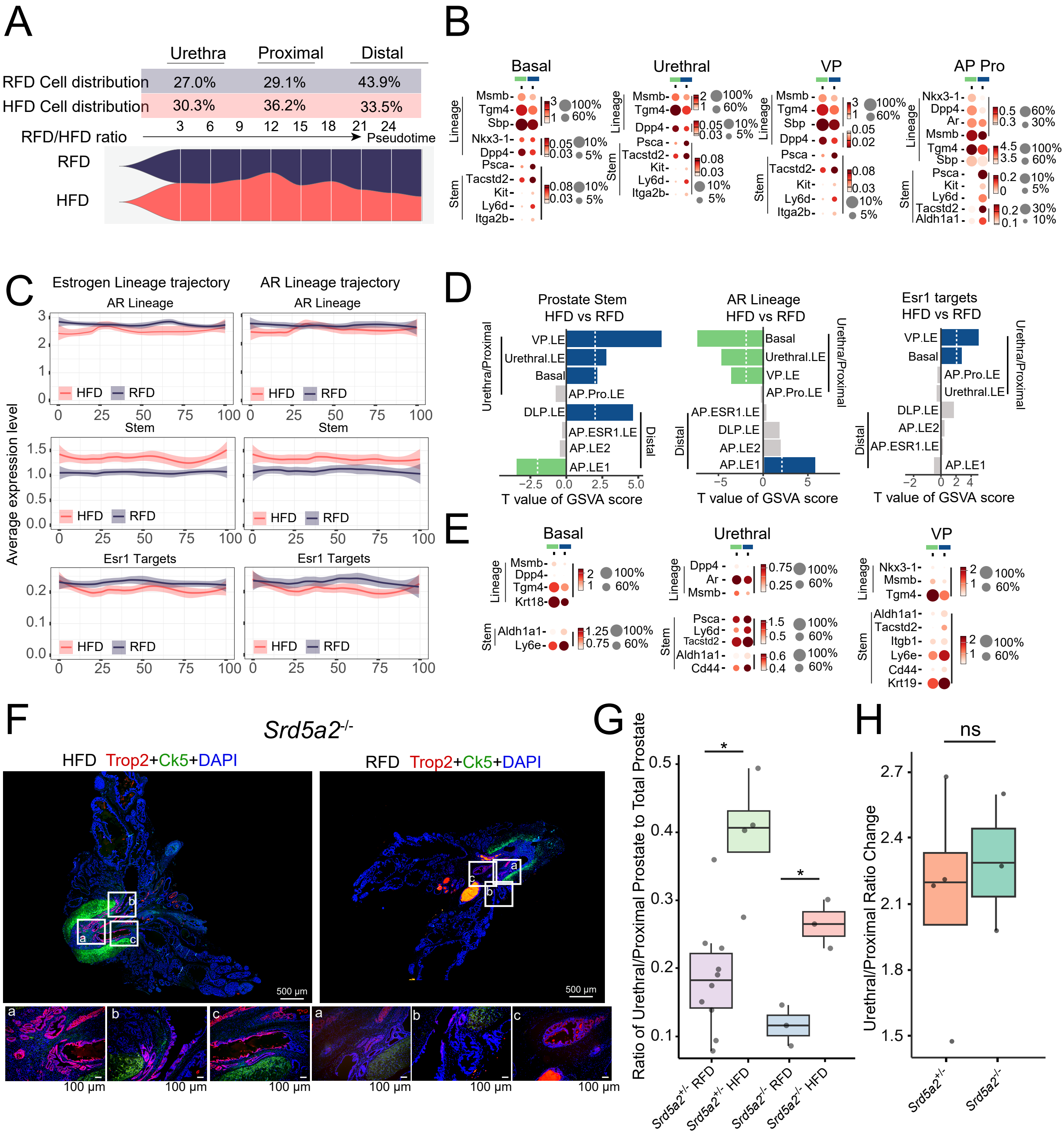

### Supplemental Figure 3

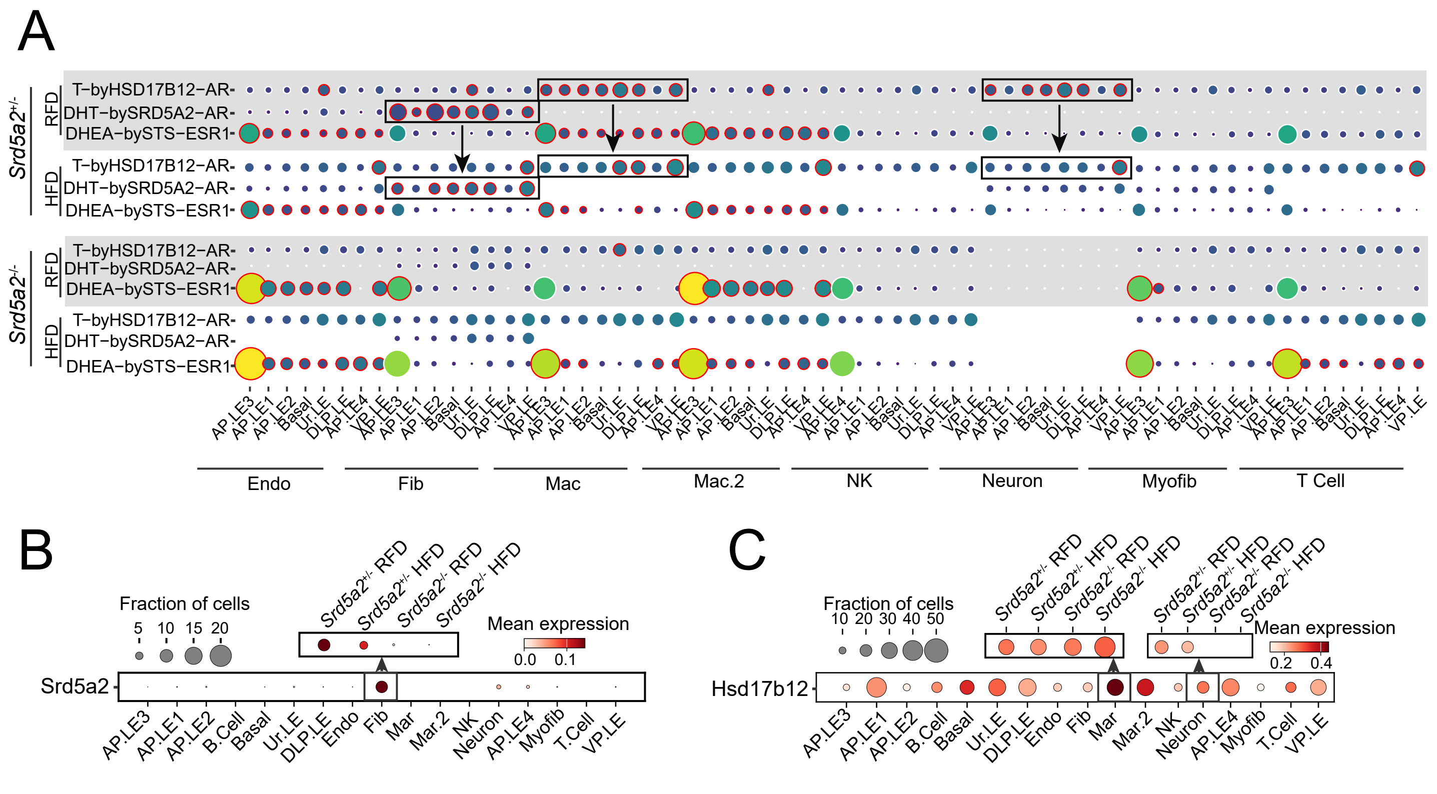

### Supplemental Figure 4

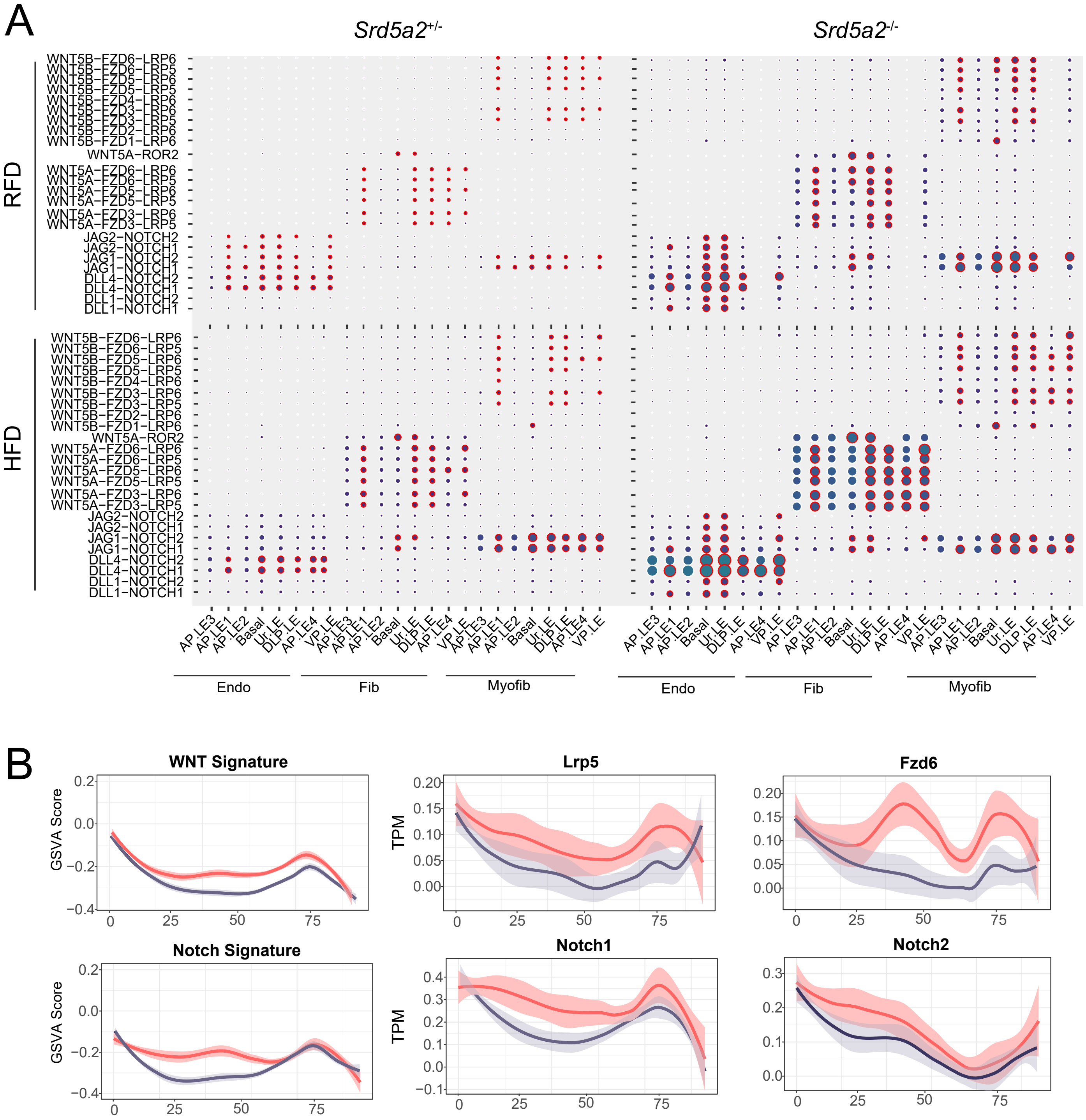

### Supplemental Figure 6

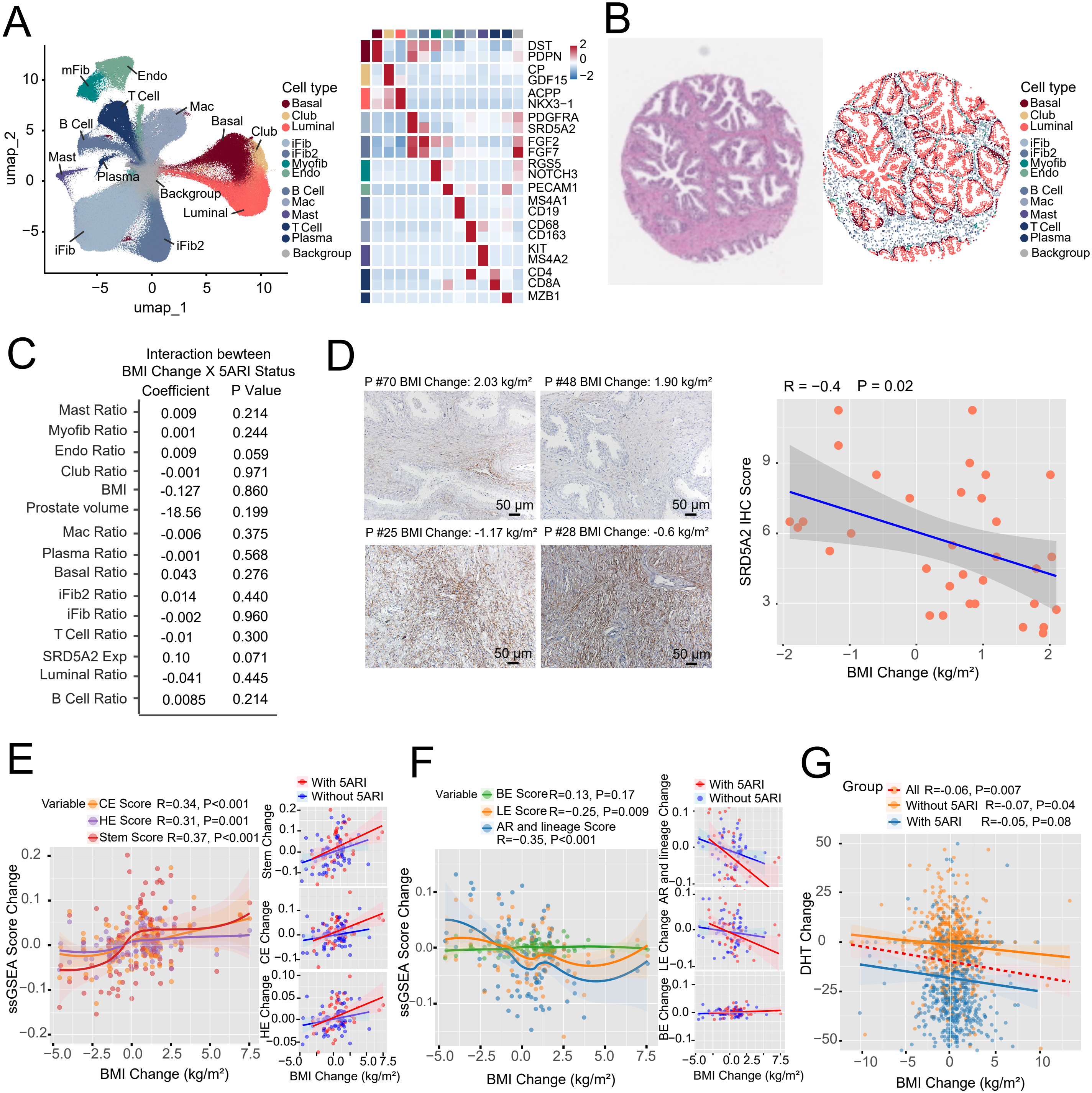

### Supplemental Figure 7

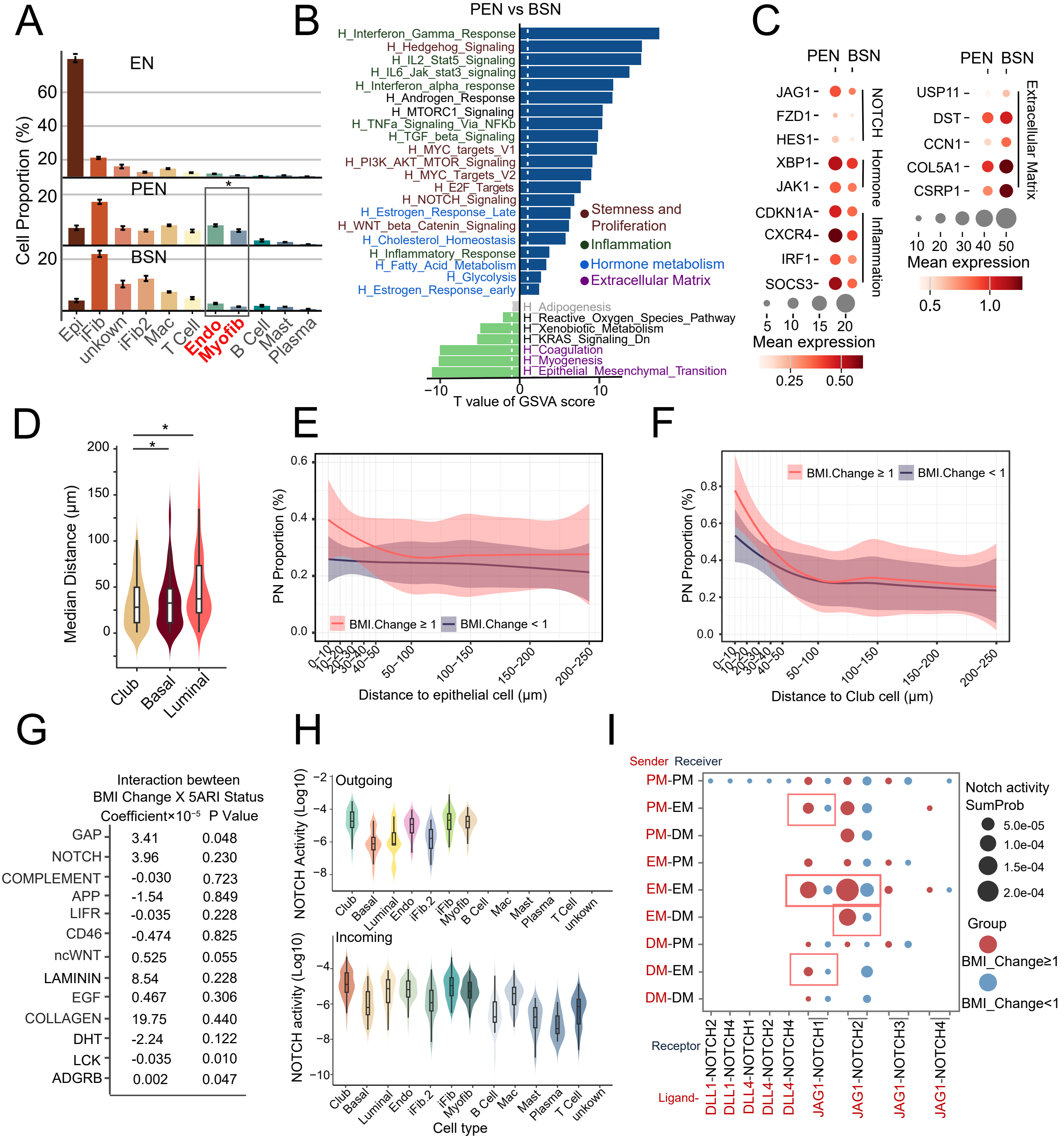

### Supplemental Figure 9

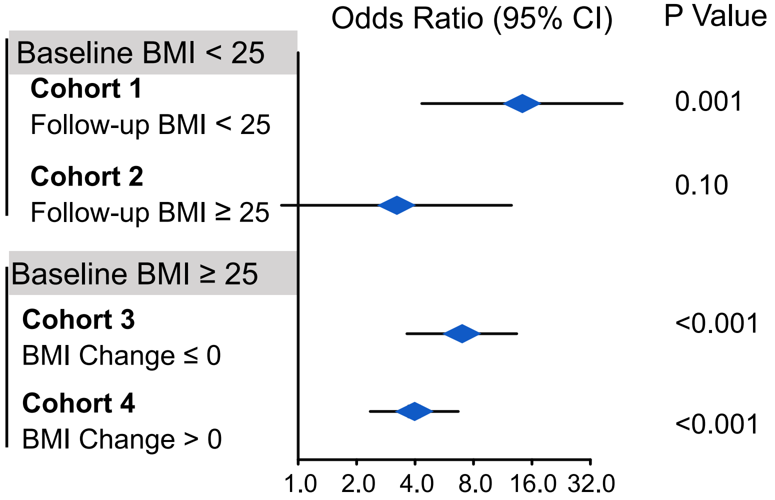
