## Supplemental Figure 5 for "Stem-like Prostate Remodeling in Obesity Mediates Resistance to 5α-Reductase Inhibition Therapy in BPH"

A Univariate Linear Regression Analyses on Factors Predicting the Growth Total Prostate Volume Growth Rate

| Variables | Regression Coefficient X 100 (95% CI) | P Value |
|---|---|---|
| 5ARI (With vs Without) | -30.66 (-34.16, -27.17) | **<0.001** |
| α-blocker (With vs Without) | 2.66 (-1.0, 6.35) | 0.16 |
| BMI Change (Continuous) | 0.95 (0.03, 1.88) | **0.044** |
| Baseline BMI (Continuous) | -0.12 (-0.55, 0.31) | 0.58 |
| Age (Continuous) | -0.27 (-0.52, -0.01) | **0.039** |
| TZI (%, Continuous) | 19.71(7.41, 32.00) | **0.002** |
| AUA Symptom Score (Continuous) | 0.01(-0.31, 0.32) | 0.96 |

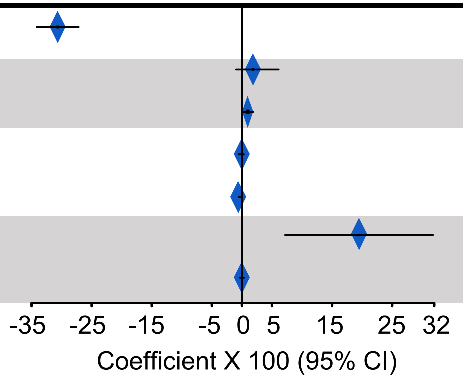


B Univariate Linear Regression Analyses on Factors Predicting the Prostate Transition Zone Volume Growth Rate

| Variables | Regression Coefficient X 100 (95% CI) | P Value |
|---|---|---|
| 5ARI (With vs Without) | -35.95 (-41.87, -30.04) | **<0.001** |
| α-blocker (With vs Without) | 2.96 (-3.12, 9.03) | 0.34 |
| BMI Change (Continuous) | 1.21 (0.03, 2.74) | 0.12 |
| Baseline BMI (Continuous) | -0.37 (-1.08, 0.33) | 0.30 |
| Age (Continuous) | -0.85 (-1.27, -0.44) | **<0.001** |
| TZI (%, Continuous) | -1.72(-1.91, -1.53) | **<0.001** |
| AUA Symptom Score (Continuous) | 0.05(-0.47, 0.57) | 0.85 |

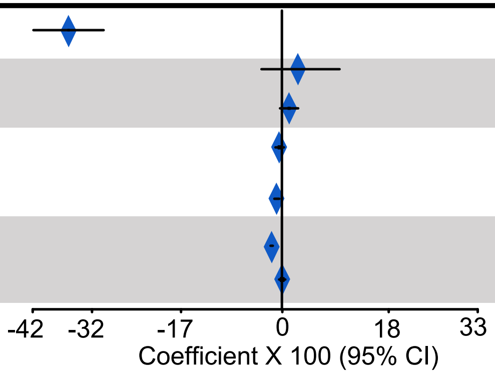


C Univariate Linear Regression Analyses on Factors Predicting the Prostate Peripheral Zone Volume Growth Rate

| Variables | Regression Coefficient X 100 (95% CI) | P Value |
|---|---|---|
| 5ARI (With vs Without) | -21.65 (-32.04, -11.26) | **<0.001** |
| α-blocker (With vs Without) | -2.39 (-12.82, 8.03) | 0.65 |
| BMI Change (Continuous) | 0.66 (-1.96, 3.29) | 0.62 |
| Baseline BMI (Continuous) | -0.61 (-1.82, 0.60) | 0.33 |
| Age (Continuous) | 0.72 (0.009, 1.44) | **0.047** |
| TZI (%, Continuous) | 2.46(2.13, 2.80) | **<0.001** |
| AUA Symptom Score (Continuous) | 0.03(-0.86, 0.93) | 0.94 |

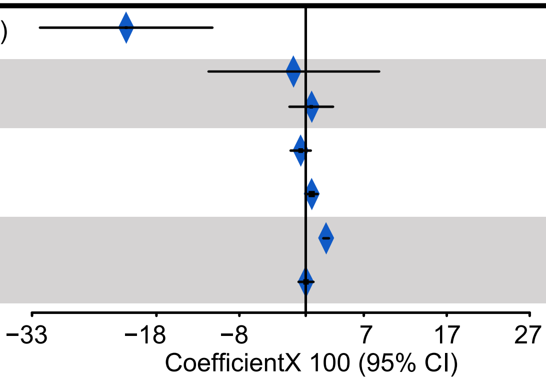
