## Supplemental Figure 8 for "Stem-like Prostate Remodeling in Obesity Mediates Resistance to 5α-Reductase Inhibition Therapy in BPH"

**A**

Multivariate Linear Regression Analyses on Factors Predicting the AUA Symptom Score Change

| Variables | Regression Coefficient (95% CI) | P Value |
|---|---|---|
| 5ARI (With vs Without) | −0.89 (−1.31, −0.47) | **<0.001** |
| α-blocker (With vs Without) | −1.34 (−1.76, −0.92) | **<0.001** |
| BMI Change (Continuous) | −0.10 (−0.21, 0.01) | 0.070 |
| Baseline BMI (Continuous) | −0.05 (−0.10, 0.00) | 0.073 |
| Age (Continuous) | 0.06 (0.03, 0.09) | **<0.001** |
| TZI (%, Continuous) | 0.64 (−0.78, 2.07) | 0.376 |
| AUA Symptom Score (Continuous) | −0.48 (−0.52, −0.45) | **<0.001** |

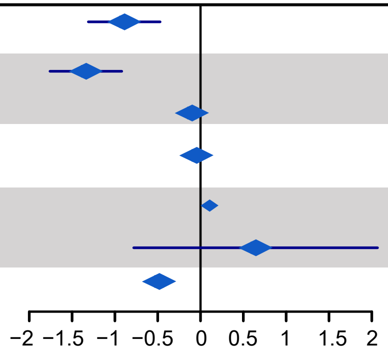


**B**

Subgroup Multivariate Linear Regression Analyses on Factors Predicting the AUA Symptom Score Change

| **Variable** | Regression Coefficient (95% CI) | P Value |
|---|---|---|
| **High-BMI-Increase subgroup** (> 2.48,n=248) | | |
| 5ARI (With vs Without) | −1.04 (−2.55 to 0.47) | 0.176 |
| α-blocker (With vs Without) | −1.69 (−3.20 to −0.18) | **0.028** |
| Baseline BMI (Continuous) | −0.09 (−0.25 to 0.08) | 0.316 |
| Age (Continuous) | 0.04 (−0.07 to 0.16) | 0.468 |
| TZI (%, Continuous) | −0.51 (−5.38 to 4.36) | 0.836 |
| AUA Symptom Score (Continuous) | −0.44 (−0.56 to −0.31) | **<0.001** |

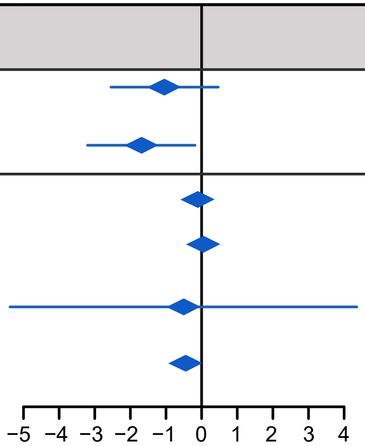


| **Variable** | Regression Coefficient (95% CI) | P Value |
|---|---|---|
| **Low-BMI-Increase subgroup** (≤ 2.48,n=2357) | | |
| 5ARI (With vs Without) | −0.89 (−1.32 to −0.45) | **<0.001** |
| α-blocker (With vs Without) | −1.31 (−1.75 to −0.87) | **<0.001** |
| Baseline BMI (Continuous) | −0.03 (−0.09 to 0.02) | 0.226 |
| Age (Continuous) | 0.07 (0.04 to 0.10) | <0.001 |
| TZI (%, Continuous) | 0.68 (−0.81 to 2.17) | 0.368 |
| AUA Symptom Score (Continuous) | −0.49 (−0.53 to −0.45) | <0.001 |

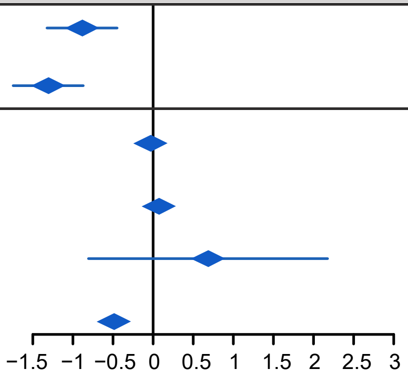
