## Supplemental Table 1 for "Stem-like Prostate Remodeling in Obesity Mediates Resistance to 5α-Reductase Inhibition Therapy in BPH"

**Table S1: Clinical demographic information of the MTOPS population (n = 2605)**

| Variable | Overall |
| --- | --- |
| Age (Mean ± SD), year | 66.97±7.30 |
| Baseline BMI (Mean ± SD), kg/m2 | 28.02±4.30 |
| **Treatment, n** |  |
| Placebo | 631 |
| Dox | 660 |
| Fin | 640 |
| Combo | 674 |
| **Initial Volume (Mean ± SD), cc** |  |
| Total prostate | 36.60±20.37 |
| Transition zone | 16.5±14.24 |
| Peripheral zone | 20.10±9.46 |
| Initial TZI | 0.41±0.15 |
| Initial AUA Score | 16.83±5.84 |
