## Supplemental Table 2 for "Stem-like Prostate Remodeling in Obesity Mediates Resistance to 5α-Reductase Inhibition Therapy in BPH"

**Table S2: Clinical demographic information of the samples used for Xenium spatial transcriptomics**

| Patient ID | Dot number | Age, year | Prostate volume, cc | BMI Change, kg/m^2^ | 5ARI use  (1: with;  0: without) | Epi Ratio,% | BMI at Surgery, kg/m^2^ |
| --- | --- | --- | --- | --- | --- | --- | --- |
| P1 | Dot3 | 70 | 37.28 | -1.68 | 0 | 0.117931547619048 | 23.73 |
| P1 | Dot1 | 70 | 37.28 | -1.68 | 0 | 0.123395336651821 | 23.73 |
| P1 | Dot2 | 70 | 37.28 | -1.68 | 0 | 0.00813008130081301 | 23.73 |
| P1 | Dot5 | 70 | 37.28 | -1.68 | 0 | 0.003094179075614 | 23.73 |
| P1 | Dot4 | 70 | 37.28 | -1.68 | 0 | 0.0180735700616628 | 23.73 |
| P11 | Dot1 | 57 | 163.02 | NA | 0 | 0.290099769762088 | 35.35 |
| P11 | Dot2 | 57 | 163.02 | NA | 0 | 0.0842062852538276 | 35.35 |
| P13 | Dot2 | 75 | 53.04 | 1.3 | 0 | 0.0623058651562214 | 44.75 |
| P13 | Dot1 | 75 | 53.04 | 1.3 | 0 | 0.664162878039138 | 44.75 |
| P15 | Dot1 | 71 | 27.04 | 0.66 | 1 | 0.532677442023893 | 25.8 |
| P15 | Dot2 | 71 | 27.04 | 0.66 | 1 | 0.412329950334701 | 25.8 |
| P15 | Dot3 | 71 | 27.04 | 0.06 | 1 | 0.0550561797752809 | 25.8 |
| P16 | Dot2 | 71 | 74.529 | 1.72 | 0 | 0.453549917868224 | 24.68 |
| P16 | Dot1 | 71 | 74.529 | 1.72 | 0 | 0.483050847457627 | 24.68 |
| P19 | Dot2 | 64 | 209.7 | NA | 0 | 0.180999839254139 | 27.88 |
| P19 | Dot1 | 64 | 209.7 | NA | 0 | 0.356118332586284 | 27.88 |
| P20 | Dot2 | 67 | 20.6 | 1.44 | 1 | 0.0586206896551724 | 27.96 |
| P20 | Dot3 | 67 | 20.6 | 1.44 | 1 | 0.417948717948718 | 27.96 |
| P20 | Dot1 | 67 | 20.6 | 1.44 | 1 | 0.00625488663017983 | 27.96 |
| P22 | Dot2 | 68 | 84.2 | -0.2 | 1 | 0.245099941781487 | 29.53 |
| P22 | Dot3 | 68 | 84.2 | -0.2 | 1 | 0.148163782186577 | 29.53 |
| P22 | Dot1 | 68 | 84.2 | -0.2 | 1 | 0.477218225419664 | 29.53 |
| P23 | Dot1 | 89 | 98.8 | -1.01 | 1 | 0.0774217447675496 | 22.81 |
| P23 | Dot2 | 89 | 98.8 | -1.01 | 1 | 0.401940545004129 | 22.81 |
| P24 | Dot2 | 74 | 37.7 | 0.3 | 0 | 0 | 20.2 |
| P24 | Dot1 | 74 | 37.7 | 0.3 | 0 | 0.513521048229718 | 20.2 |
| P25 | Dot3 | 75 | 80 | -1.17 | 0 | 0.00360824742268041 | 30.3 |
| P26 | Dot1 | 59 | NA | -0.78 | 0 | 0.626732673267327 | 19.49 |
| P26 | Dot2 | 59 | NA | -0.78 | 0 | 0.587369480437791 | 19.49 |
| P27 | Dot2 | 73 | 18.2 | -0.1 | 1 | 0.168779517772806 | 22.12 |
| P27 | Dot1 | 73 | 18.2 | -0.1 | 1 | 0.360249466601018 | 22.12 |
| P28 | Dot3 | 64 | 84.6 | -0.6 | 0 | 0.00196818107265868 | 30.7 |
| P28 | Dot2 | 64 | 84.6 | -0.6 | 0 | 0.0489838457529964 | 30.7 |
| P28 | Dot1 | 64 | 84.6 | -0.6 | 0 | 0.371494556252062 | 30.7 |
| P29 | Dot2 | 70 | 23.4 | 1.8 | 1 | 0.404754829123328 | 25.79 |
| P29 | Dot1 | 70 | 23.4 | 1.8 | 1 | 0.333333333333333 | 25.79 |
| P32 | Dot1 | 60 | 59 | 2.2 | 1 | 0.0738330341113106 | 26.08 |
| P32 | Dot2 | 60 | 59 | 2.2 | 1 | 0.280738786279683 | 26.08 |
| P33 | Dot2 | 76 | 222 | 0.84 | 1 | 0.20770986558458 | 21.76 |
| P33 | Dot1 | 76 | 222 | 0.84 | 1 | 0.151060358890701 | 21.76 |
| P34 | Dot3 | 76 | 75.8 | 2.01 | 0 | 0.00291028723269644 | 24.72 |
| P34 | Dot1 | 76 | 75.8 | 2.01 | 0 | 0.364406779661017 | 24.72 |
| P34 | Dot2 | 76 | 75.8 | 2.01 | 0 | 0.120971302428256 | 24.72 |
| P35 | Dot2 | 68 | 23.3 | 0.88 | 0 | 0.66796317904836 | 28.8 |
| P35 | Dot1 | 68 | 23.3 | 0.88 | 0 | 0.0038961038961039 | 28.8 |
| P36 | Dot1 | 64 | 117.4 | 0.21 | 0 | 0.00255156283223474 | 26.2 |
| P36 | Dot2 | 64 | 117.4 | 0.21 | 0 | 0.234925514305982 | 26.2 |
| P38 | Dot1 | 79 | 10.7 | NA | 0 | 0.00109443402126329 | 28.3 |
| P38 | Dot2 | 79 | 10.7 | NA | 0 | 0.00346565847511027 | 28.3 |
| P38 | Dot3 | 79 | 10.7 | NA | 0 | 0.00196831020568842 | 28.3 |
| P40 | Dot1 | 75 | 42.1 | -1.3 | 0 | 0.727094880991197 | 28.1 |
| P40 | Dot2 | 75 | 42.1 | -1.3 | 0 | 0.419247260600286 | 28.1 |
| P41 | Dot1 | 66 | 197.3 | 1.91 | 0 | 0.192013129102845 | 28.2 |
| P41 | Dot2 | 66 | 197.3 | 1.91 | 0 | 0.0373751783166904 | 28.2 |
| P42 | Dot1 | 63 | 79.6 | 0.2 | 1 | 0.308553728853943 | 28.05 |
| P42 | Dot3 | 63 | 79.6 | 0.2 | 1 | 0.541858678955453 | 28.05 |
| P42 | Dot2 | 63 | 79.6 | 0.2 | 1 | 0.0255564715581204 | 28.05 |
| P43 | Dot1 | 75 | 107.6 | 1.2 | 1 | 0.420221891799154 | 27.6 |
| P43 | Dot2 | 75 | 107.6 | 1.2 | 1 | 0.000271997824017408 | 27.6 |
| P43 | Dot3 | 75 | 107.6 | 1.2 | 1 | 0.00228211258422082 | 27.6 |
| P44 | Dot2 | 71 | 42.1 | -0.98 | 1 | 0.440803070563921 | 21.1 |
| P44 | Dot1 | 71 | 42.1 | -0.98 | 1 | 0.475046010609505 | 21.1 |
| P45 | Dot2 | 82 | 32.2 | -1.78 | 1 | 0.0747046560111188 | 24.28 |
| P45 | Dot1 | 82 | 32.2 | -1.78 | 1 | 0.00874125874125874 | 24.28 |
| P46 | Dot2 | 73 | 21 | 0.9 | 0 | 0.398277172536103 | 26.1 |
| P46 | Dot1 | 73 | 21 | 0.9 | 0 | 0.00860585197934595 | 26.1 |
| P47 | Dot1 | 70 | 40.9 | 0.7 | 0 | 0.544516829533116 | 28.5 |
| P47 | Dot1 | 70 | 40.9 | 0.7 | 0 | 0.0302937576499388 | 28.5 |
| P48 | Dot1 | 49 | 36.9 | 1.9 | 1 | 0.350923872875092 | 27.5 |
| P48 | Dot2 | 49 | 36.9 | 1.9 | 1 | 0.495738636363636 | 27.5 |
| P49 | Dot1 | 73 | 28 | 0.5 | 0 | 0.49044406970208 | 30.7 |
| P49 | Dot2 | 73 | 28 | 0.5 | 0 | 0.00306211723534558 | 30.7 |
| P5 | Dot1 | 88 | 101.92 | 2.21 | 0 | 0.270353302611367 | 28.59 |
| P5 | Dot1 | 88 | 101.92 | 2.21 | 0 | 0.0306709265175719 | 28.59 |
| P52 | Dot2 | 67 | 29.1 | 1.04 | 0 | 0.405939123979213 | 30.05 |
| P52 | Dot1 | 67 | 29.1 | 1.04 | 0 | 0.248399487836108 | 30.05 |
| P53 | Dot2 | 56 | 276.8 | 0.67 | 0 | 0.56250657133845 | 25.8 |
| P53 | Dot1 | 56 | 276.8 | 0.67 | 0 | 0.493252183117227 | 25.8 |
| P54 | Dot1 | 72 | 187.2 | 1.85 | 0 | 0.351885098743268 | 21.3 |
| P54 | Dot2 | 72 | 187.2 | 1.85 | 0 | 0.161845882093177 | 21.3 |
| P54 | Dot3 | 72 | 187.2 | 1.85 | 0 | 0.119449656035022 | 21.3 |
| P58 | Dot3 | 77 | 86 | 0.8 | 0 | 0.000963921784632333 | 27.2 |
| P58 | Dot1 | 77 | 86 | 0.8 | 0 | 0.0828670959569302 | 27.2 |
| P58 | Dot2 | 77 | 86 | 0.8 | 0 | 0.408918005071851 | 27.2 |
| P59 | Dot2 | 60 | 252.8 | 2.1 | 0 | 0.00139567341242149 | 26.8 |
| P59 | Dot1 | 60 | 252.8 | 2.1 | 0 | 0.266615027110767 | 26.8 |
| P6 | Dot2 | 65 | 57.408 | 1.9 | NA | 0.352597602213342 | 27.95 |
| P6 | Dot3 | 65 | 57.408 | 1.9 | NA | 0.416448906615321 | 27.95 |
| P6 | Dot1 | 65 | 57.408 | 1.9 | NA | 0.206715210355987 | 27.95 |
| P60 | Dot1 | 68 | 69.5 | 0.98 | 0 | 0.714790509843513 | 27.3 |
| P60 | Dot2 | 68 | 69.5 | 0.98 | 0 | 0.602336274387971 | 27.3 |
| P60 | Dot3 | 68 | 69.5 | 0.98 | 0 | 0.198626447616483 | 27.3 |
| P62 | Dot2 | 69 | 183.2 | -1.7 | 0 | 0.000853242320819113 | 25.3 |
| P62 | Dot1 | 69 | 183.2 | -1.7 | 0 | 0.145757071547421 | 25.3 |
| P63 | Dot1 | 67 | 72.9 | NA | 1 | 0.00231481481481481 | NA |
| P65 | Dot2 | 67 | 170.4 | 1.2 | 0 | 0.384898710865562 | 28.3 |
| P65 | Dot1 | 67 | 170.4 | 1.2 | 0 | 0.440511307767945 | 28.3 |
| P66 | Dot1 | 65 | 130.6 | -1.9 | 0 | 0.0009000900090009 | 23.9 |
| P66 | Dot3 | 65 | 130.6 | -1.9 | 0 | 0.000440625688477638 | 23.9 |
| P67 | Dot1 | 68 | 49.7 | 1.6 | 0 | 0.184760865611929 | 26.6 |
| P67 | Dot2 | 68 | 49.7 | 1.6 | 0 | 0.342682725622835 | 26.6 |
| P68 | Dot1 | 53 | 18.2 | 0.4 | 1 | 0.0813526096544964 | 30.8 |
| P68 | Dot2 | 53 | 18.2 | 0.4 | 1 | 0.314212548015365 | 30.8 |
| P69 | Dot2 | 58 | 171.5 | 0.8 | 0 | 0.0121278941565601 | 23.1 |
| P69 | Dot3 | 58 | 171.5 | 0.8 | 0 | 0.259019741320626 | 23.1 |
| P70 | Dot1 | 74 | 91.2 | 2.03 | 0 | 0.308917197452229 | 26.2 |
| P70 | Dot2 | 74 | 91.2 | 2.03 | 0 | 0.00120789155818011 | 26.2 |
| P72 | Dot3 | 67 | 173 | 1.8 | 0 | 0.0155223880597015 | 37.3 |
| P73 | Dot2 | 68 | 119.9 | 1.77 | 0 | 0.0050125313283208 | 28.5 |
| P73 | Dot1 | 68 | 119.9 | 1.77 | 0 | 0.31986531986532 | 28.5 |
| P74 | Dot1 | 62 | 63.9 | -2.73 | 0 | 0.00744532340623546 | 23.9 |
| P74 | Dot2 | 62 | 63.9 | -2.73 | 0 | 0.0119165839126117 | 23.9 |
| P74 | Dot3 | 62 | 63.9 | -2.73 | 0 | 0.0197830248883216 | 23.9 |
| P75 | Dot2 | 78 | 122.7 | 0.15 | 1 | 0.280330482678649 | 23.78 |
| P75 | Dot1 | 78 | 122.7 | 0.15 | 1 | 0.501251042535446 | 23.78 |
| P76 | Dot3 | 67 | 45 | -1.17 | 1 | 0.00334378265412748 | 29.8 |
| P76 | Dot1 | 67 | 45 | -1.17 | 1 | 0.00237944162436548 | 29.8 |
| P76 | Dot2 | 67 | 45 | -1.17 | 1 | 0.00148809523809524 | 29.8 |
| P78 | Dot2 | 62 | 62.5 | -0.28 | 0 | 0.429469375822248 | 27.12 |
| P79 | Dot2 | 82 | 207 | 0.54 | 0 | 0.334795963141729 | 26.3 |
| P79 | Dot1 | 82 | 207 | 0.54 | 0 | 0.031055900621118 | 26.3 |
| P80 | Dot2 | 66 | 16 | NA | 1 | 0.000309405940594059 | NA |
| P80 | Dot1 | 66 | 16 | NA | 1 | 0.0024390243902439 | NA |
| P80 | Dot3 | 66 | 16 | NA | 1 | 0.00440426518312471 | NA |
| P81 | Dot2 | 71 | 78 | 1 | 0 | 0 | 32 |
| P81 | Dot1 | 71 | 78 | 1 | 0 | 0.352212069121505 | 32 |
| P81 | Dot3 | 71 | 78 | 1 | 0 | 0.00914526908195568 | 32 |
| P82 | Dot3 | 63 | 107.1 | 0.8 | 0 | 0.0142899672521584 | 29.41 |
| P82 | Dot1 | 63 | 107.1 | 0.8 | 0 | 0.00114126299771747 | 29.41 |
| P82 | Dot2 | 63 | 107.1 | 0.8 | 0 | 0.00164925783397471 | 29.41 |
| P83 | Dot1 | 72 | 106.6 | 0.83 | 0 | 0.00530035335689046 | 21.94 |
| P83 | Dot2 | 72 | 106.6 | 0.83 | 0 | 0.00112497443239926 | 21.94 |
| P85 | Dot3 | 62 | 33.7 | 1.38 | 1 | 0.00136379133992499 | 28.58 |
| P85 | Dot1 | 62 | 33.7 | 1.38 | 1 | 0.00158084914182475 | 28.58 |
| P85 | Dot2 | 62 | 33.7 | 1.38 | 1 | 0.0295774647887324 | 28.58 |
| P86 | Dot2 | 68 | 50.4 | 0.89 | 0 | 0.302516411378556 | 30.1 |
| P86 | Dot1 | 68 | 50.4 | 0.89 | 0 | 0.214427719397898 | 30.1 |
| P87 | Dot2 | 77 | NA | 0.8 | 1 | 0.00355450236966825 | 29.4 |
| P87 | Dot1 | 77 | NA | 0.8 | 1 | 0.0350404312668464 | 29.4 |
| P88 | Dot1 | 75 | 180 | 0.87 | 0 | 0.129610655737705 | 24.6 |
| P88 | Dot2 | 75 | 180 | 0.87 | 0 | 0.498877917414722 | 24.6 |
| P89 | Dot1 | 72 | 99.3 | 1.3 | 0 | 0.183501683501683 | 30.4 |

NA: Not Available; cc: cubic centimeter
