## Supplemental Table 3 for "Stem-like Prostate Remodeling in Obesity Mediates Resistance to 5α-Reductase Inhibition Therapy in BPH"

**Table S3: Gene sets used in the article**

| Prostate Stem Marker | Concluded from reference 1 to 6. |
| --- | --- |
|  | Tacstd2 Itgb1 Cd44 Krt4 Krt19 Itga2b Sox2 Prom1 Ly6d Ly6e Aldh1a1 Psca Alcam Kit |
| LE Signature | Calculated from reference 6 |
|  | Npy Acpp Msmb Klk1b8 Rdh11 Nefh Nkx3-1 Dhrs7 Mme Cpe Pak1ip1 Dbi Ncapd3 Tff3 Pdlim5 Fads2 Azgp1 Steap2 Sec11c Dpp4 Klk4 Pdia3 Mia3 Sord Sms Slc45a3 Abcc4 Tmprss2 Slc39a6 Lrrc26 Naaa Hsp90b1 Tmeff2 H2afj Tspan1 Aplp2 Asrgl1 Mesp1 Spon2 Pmepa1 Acsl3 Slc39a10 Iqgap2 Dcxr Rab3b Slc30a4 Glo1 Vgll3 Spdef Idh1 Alox8 Erlec1 Fkbp5 Slc4a4 Bnip3 Ncoa4 Golgb1 Rab27a Arg2 Clgn Nucb2 Amd1 Nipal3 Ppp3ca Tpd52 Rpn2 Dnajc3 Arfgap3 Rcan3 Spock1 Zcchc6 Scd1 Acat3 Lcp1 Ergic1 Slc2a12 Abhd2 Lman1 5330417C22Rik Zfp51 Golm1 Galnt7 Ctbs Adi1 Mpc2 Bloc1s6 Aldh6a1 Rbm47 Mgst1 Lsamp Steap1 Utrn Creb3l4 Gusb Atp6v1g1 Atp2c1 Antxr2 Mboat2 Ckb Canx Tmem141 Gcc2 Gadd45g Ramp1 Fam213a Glud1 Acsm3 Bcas1 Arhgap6 Anpep Homer2 Gmpr Uqcrq Tm9sf3 Entpd5 Nme4 Tmem192 Bmpr1b Aff3 Nudt4 Syap1 Txndc16 Nans Alcam Mcfd2 Cd46 Sec63 Mccc2 Pebp4 Copb1 Gcnt2 Pcyox1 Smim14 Slc12a2 Casc4 Arl6ip5 Aldh9a1 Ankrd37 Syt7 Pgm3 Fasn Kif5c Hoxb13 Gtf3c6 Tax1bp1 Taok3 Degs1 P4hb Tmem87a Endod1 Golga4 Glrx2 Hist1h2ao Trpm8 Psma7 Ubxn4 Hipk2 Ssr3 Fam3b Minos1 Soat1 Ptprf Agtrap Ttc3 Ssr4 Tstd1 Elovl5 Rab2a Ppm1k Ddt Lamp2 Gfpt1 Slc44a4 Thoc2 Ppcs Dnajc10 Basp1 Tcea3 Lifr Thumpd1 Hebp2 Tmbim6 Tusc3 Tmem79 Psma4 Kif21a Hist1h1c Syngr2 Cmas Fxyd3 Ufl1 Dhcr24 Atp6v0b Npnt Herpud1 Tspan8 Pebp1 Ephx2 Kif5b Capzb Fkbp3 Tmco1 Mien1 Flnb 1110004F10Rik Ttc37 Rab27b Ebp Tars Sec23b Pdxdc1 Ptov1 Scp2 Epcam Map7 Ebag9 Mlec Rnf13 Xbp1 Maged2 Uqcr10 Coprs Maf Npdc1 Steap4 Eif5b Foxa1 Fbxo25 Trim24 Ccng2 Dsc2 Mgst3 Ube2j1 Svip Leo1 Ar Psap Itm2c Cyb561 R3hdm2 Cd2ap Gpx1 Cbr1 Rb1cc1 Mrps35 Acsl1 Ndufa3 Acbd5 Meaf6 Cyb5 Zfhx3 Sec62 Sec61b Impact Mzt2 4931406C07Rik Rad21 Fam210b AA986860 Ndufa4 Snx4 H1f0 Serf2 Ssr1 Ppdpf Atp6v1d Atp6ap1 Cndp2 Suclg1 Pdia4 Tmc5 Rhobtb3 1700025G04Rik Zkscan1 Atrx Tsc22d3 Ndrg1 Sc5d Mlph Secisbp2l Aldh1a3 Fam129a Xiap Cox7a2 Ang Ppp1r7 Trip11 Tbx3 H13 Zcrb1 Ndufa13 Slc31a1 Tmem230 Prdx4 Zbtb20 Romo1 Sel1l Cox17 Cpne3 Srp72 Psma2 Usp14 Eid1 Pdia6 Prpf40a Wwc1 Atp8b1 Limch1 Ap2s1 Spcs2 Nbr1 Copb2 Serpinb6a Larp7 Mtdh Nt5dc1 Armcx3 Anxa3 Coa3 Dcaf6 Rdx Tmed4 Snrpn Tmx4 Thrap3 Bri3 Aldh3a2 Slc39a7 Tmed10 Smarca1 Emc7 Ndufa2 Kmt2c Daam1 Vamp2 Acat1 Il1r1 Mrp63 Ociad1 Tram1 Prdx3 Adh5 Pnkd Uqcr11 Eif2ak4 Aldh7a1 Anapc11 Cul5 Ssb Itsn2 Col4a3bp Kctd3 Nedd4l Tmbim4 Tm9sf2 Cdc42ep5 4933434E20Rik Zfml Tbca Idh2 Prss8 Cst3 G3bp2 Reep5 Smarca5 Psmd7 Trove2 Mrpl41 Nars Sar1b Cd9 Spcs1 Lypla1 Bcap29 Dsg2 Atf6 Insig1 Nucks1 Arhgap29 Snrnp27 Sec61g Cat Usp8 Cobll1 Idi1 Magt1 Ergic2 Dhx29 Atxn7l3b Ormdl2 Ndufc1 Ndufv3 Spcs3 LOC102631912 U2surp Eif2ak2 Tbc1d15 Fam120a Phpt1 Nsmce1 Ccdc47 Tmed2 Ndufa5 Calu Pcmtd1 Lrrfip1 Pja2 BC005624 Ralbp1 Cltc Eif3a Nop10 Eea1 Rps26 Zfp106 Gtf2h5 Rfc1 Rab4a Vdac1 Tmem179b Hexb Hdlbp Cnpy2 Ifnar1 Tcea1 Cep350 Tbl1xr1 Fkbp2 Sft2d1 Scoc Phax Nupr1 Rps27l Tmem14c Echs1 Cox6a1 Srp14 Ptgr1 Mtif3 St6gal1 Galnt1 Smarcc1 Plekhb2 Gabarapl2 Ofd1 Golph3 Dstn Tmem256 Dnajc1 Bola3 Scarb2 Calr Prdx6 Eif2ak1 Tmem258 Tmf1 Phf3 Hsd17b4 Hp1bp3 Hspa9 Kdelr2 Rpl23 Fam192a B2m Cmtm6 Golga2 Ndufb9 Tmed3 Fdft1 Dynlrb1 Vegfa Cwc15 Sec31a Ndfip1 Lyrm2 Os9 Leprot Tmed9 Bccip Rpn1 Usp16 Pdcd4 Nbl1 Mrpl33 Tceal8 Vps36 Tmem50a Vps35 Erbb3 Grb2 Acaa1a Prdx2 Dpy30 Mrpl36 Zbtb38 Ltf Pcbd1 Smdt1 Akr1a1 Ndufa11 Gnl3 Rab13 Ube2l3 C1d Chmp2b Uso1 Txndc17 Dnajc19 Senp6 Baz1b Clip1 Ubtf Psmb5 Serinc1 Txnl4a Tpr Erp29 Bnip3l Tmem134 Rpl31 Vcp Usp7 Cox8a Ndufc2 Dsp Hspe1 Ift57 Zfr Ttc1 Yme1l1 |
| HE Signature | Calculated from reference 6 |
|  | Krt13 Krt17 Krt15 Mmp7 Krt19 Ly6d Krt5 S100a6 Krt7 Krt23 Cldn4 Hbegf Plat Tm4sf1 Wfdc2 Tacstd2 Lamb3 S100a9 Olfm4 Slpi Gadd45a Klf5 Cldn1 Aqp3 Sfn Anxa2 G0s2 Elf3 Gstp1 Actg1 Plau Tnfrsf12a Phlda1 Ly6e Hspb1 Plcg2 Gprc5a Itga2 Fhl2 Igfbp3 S100a14 Areg Gm5068 Phlda2 Gpx2 Pdlim1 Perp Zfp36l1 Rarres1 Crabp2 F3 Csta Hmga1-rs1 Mpzl2 Gabrp Plaur 1110008P14Rik Slc14a1 Niacr1 Btg3 Cxcl17 Ass1 Sdc1 Asph S100a16 Ppp1r15a Tnfaip2 Ets2 Trim29 Ybx3 Meis2 Vmp1 Sgk1 Actb Plin2 Dusp5 Errfi1 Lypd3 Mast4 Atp1b1 Adm Serpinb5 Mgp Ddit4 Ctsb Sdc4 Lima1 Glul Eef1b2 Pitx1 Cyp4b1 Rap2b Cldn7 Rps18 Rps2 Lmna Krt8 Cd82 Rps19 Tmprss4 Sqstm1 Phlda3 Ubc Serpinb1a Id1 Clu Rpl10a Rps9 Foxq1 Rab11fip1 Fam110c Epha2 Pkm Anxa1 Ifi204 Prss22 Fosl1 Rps3 Rps14 Jup Rhov Inpp1 Rpl13 Ier3 Gm6404 Cav2 Myc Tuba1c Rnd3 Ccnd1 Cd59b Chchd10 Btf3 Mdk Cyr61 Ezr Tes Dsc3 Ctsh Bag1 Atf3 Dennd2c Itpkc Itga6 Rps21 Irf6 Lcn2 Rps8 Taf1d Rpl35 Tkt Gpr87 Rel Maff Sh3bgrl3 Luzp1 Pcdh7 Syne2 Fam3c Ptpn2 Rpl3 Nfkbia Kcnk1 Tmem40 Irf2bp2 Bhlhe40 Hist2h4 Myof Plp2 Nabp1 Trip6 Gm5506 Arpc2 Tubb4b Ncoa7 Dhrs3 Rpl18a Rpl7a Ppa1 Rpl37a Eif4a1 Cd44 Cebpb Rpl37 Ddx21 Rpl21 Tagln2 Actn4 Jun Rgs12 Rpl8 Ntn4 Ralgds Prdx5 Gm6251 Rpl24 Zfand5 Gm4604 Slc2a1 Rps7 Lgals3 Cited4 Wee1 Slc25a5 Rpsa Rps23 Ier5 Rplp0 S100a13 Rpl27a Stom Sri 2010111I01Rik Rpl11 Naca Rpl14 Rpl19 Rpl12 Rps11 Frmd6 Fbl Rps6 Rps12 Tgif1 Pdlim4 Tnfrsf21 Cd55 Dnajb6 Rps15a Rpl26 Rpl28 Rpl15 Rpl9 Rpl4 Eif5 Bcl10 Rps27a Ctss Btg1 Cdkn1a Lad1 Gna15 Ankrd11 Ddx5 Rpl35a Palld Rpl30 Rpl7 Rps29 Gapdh Higd2a Ehf Rpl10 Cycs Hes1 Rpl39 Polr1d Rps28 Mdm2 Rps17 Rps5 Jund Scpep1 Mafb Gbp2 Eef1d Ctsd Efhd2 Rps24 Coro1c Lrrc8a Gltp Ceacam1 Rpl22 Jag1 Rtn4 Krt18 H3f3b Ehd4 Sema4b Prmt1 Rnf145 Pptc7 Optn Ifitm3 Ugcg Rps25 Fau Map2k3 Midn Eef1a1 Capn1 Traf4 Cnn2 Tle4 Ahr Srsf7 Lmo4 Itgb1 Tra2b Ppp1r14b Zswim6 Ints6 Btg2 Ddr1 Oser1 Srpk1 Npm1 Arl4a Vps37b Rpl13a Rpl5 Pim1 Snrpf Ptpn13 Slfn5 Gja1 Sod2 Vdac2 Akirin2 Hnrnpab Tsc22d2 Igfbp2 Pim3 Kdsr Slc38a2 Spint1 Rpl38 Pnn Klf6 Drap1 Rpl6 Cxcl16 Agr2 Gpc1 Rps20 Ube2s Baiap2 Aldh2 Rps16 Zfp36l2 Gm9763 Chpt1 Capg Snrpb Crot Eif6 Gm7866 Shb Eif3e Rap1b Tpt1 Ldha Rab10 Cltb Nfe2l2 Camta1 Arl6ip4 Baz1a Pa2g4 Aes Sun1 F11r Pbx1 Fosb |
| CE Signature | Calculated from reference 6 |
|  | Scgb3a1 Scgb1a1 Olfm4 Pigr Slpi Wfdc2 Rarres1 Mgp Cp Lcn2 Cxcl17 Atp1b1 Mmp7 Elf3 Crabp2 Tff1 Ly6e Zfp36l1 Krt7 Psca Scube2 Krt19 Tnfaip2 S100a6 Cd82 Cldn4 Agr2 Hes1 Rhov Klf5 Clu Lxn Slc40a1 Igfbp3 Tspan3 Ltf Meis2 Oit1 Ass1 Gstp1 Cldn3 Lgals3 Jun Cd74 Tmem45b F3 Selenbp1 Gprc5a Chpt1 Robo1 Gnptab Gdf15 Id1 Ppp1r1b Fos Pdzk1ip1 Syne2 Nfkbiz Hist2h4 Asph Cldn7 Ncoa7 Ccnd1 Btg3 Creb3l1 Ctss Odf2l Ephx1 Sytl2 S100a13 Tnfsf10 Sdc4 Serpinb1a Slc25a29 Rgcc Ascc3 Npc2 Ctsb Sri Gm5068 Lima1 Bag1 Cyba Tmem63a Hnmt Ctsh Tnfrsf21 Cebpd Mpzl2 Gata3 Prss22 Lamb3 Anxa2 Ptprk Atf3 Cd55 Sorbs2 Ehf Rab11fip1 Thsd4 Rnd3 Vamp8 Slfn5 Slc14a1 Rnf145 Mdk Pnpla2 Liph S100a14 Ier2 Sox9 Vmp1 Krt8 Insr Fnbp4 Tspan6 Cxcl1 Klk11 Glul Add3 Hsd17b11 Wsb1 Gm11127 Shroom1 Spint2 Ypel3 Crym Tspan15 Scnn1a S100a16 Erbb2 Ccnl1 Ttll7 Dhrs3 Jup Vsig2 Birc3 Gstk1 Enah Rbpms Optn Tob1 Etl4 Spint1 Aqp3 Stt3b Limch1 Tsc22d1 Pim3 Bhlhe40 Cnfn Capn2 Tes Cxadr Papss1 Baz2b Krtcap2 Map3k5 Fhl2 H2-DMa Pbx1 Tacstd2 Anxa11 1110008P14Rik Chd9 Nfkbia Krt18 Hook2 Arhgdib Oat Fam3c Rnf213 Gltp Smim22 Tmem238 Higd1a Tjp1 Sorl1 Dusp1 H2-K1 Fbp1 Laptm4a Arglu1 Ddr1 Myof Sh3glb2 Pdlim4 Txnip Lmo7 St6gal1 Fcgrt Sf3b1 Tra2b Rsbn1l |
| BE Signature | Calculated from reference 6 |
|  | Igfbp7 Gm13889 A2m Sparcl1 Cfd Vim Tagln Tm4sf1 Myh11 Ccl12 Ier3 Socs3 Myl9 Nfkbia Krt17 Pdlim1 Gadd45a Lmna Junb H2-Ea-ps S100a6 Ppp1r15a Ybx3 Cxcl1 Krt15 G0s2 Mgp Egr1 Ubc Fosb Ets2 Rps9 Fos Hspa1a Jund Nr4a1 Emp1 Zfp36 Krt19 Actg1 Ldha Gm6404 Ifitm2 Rpl3 Cebpb Cdkn1a Rps18 Actb Eef1b2 Rpl35 Eif4a1 Cd74 Dnajb1 Rpl10a Rps2 Gm6251 Tagln2 Rps11 Rpl7 Atf3 Btg2 Rps3 Rpl38 Krt13 Rpl10 Clu Rps23 Tpt1 Rpl13 Rpl37a H3f3b Naca Atp1b3 Rps12 Rps14 Id3 Rps15a Rpl30 Sod2 Zfp36l1 Rpl9 Ier2 Rpl7a Tuba1b H2afz Gm7866 Rpl24 Rpl19 Sdcbp Gadd45b Ddit4 Mcl1 Rpl11 Rpsa Rpl18a Rps27a Srsf3 Jun Rpl39 Rpl28 Cebpd Rps7 Btf3 Rpl22 Eef1a1 Btg1 Rps25 Zfp36l2 Id1 Pfdn5 H2-K1 Ddx21 Rpl37 Rpl35a Gm4604 Rps29 Gapdh Tomm7 Cd59b Hspa8 Rps28 Rps8 Rpl4 Timp1 Rpl26 Eef1d Rps19 Pnrc1 Snrpb Rps24 Gm5506 Klf6 Rps20 Rpl6 Fau Anxa2 Rpl21 Sfn Oaz1 Gstp1 Rpl8 Phlda2 Cycs Rpl12 Ftl1 Atf4 Ube2d3 Rpl15 H2-T23 Dusp1 Rps4x Ddx5 Srsf2 Myl12b Ifitm3 Rps6 Eif3f Eef2 Pim3 Hnrnph1 Ppa1 Eif3e Rpl13a Sh3bgrl3 Dnaja1 Myl6 Nfe2l2 Rps21 Brd2 Rpl5 Tubb4b Rpl27a Anxa1 Cd81 Srsf7 Npm1 Ybx1 Hnrnpdl Aqp3 Glul Ccnl1 Rps5 Ran Elf3 Rps13 Cldn4 Vdac2 Eif5 Gm9769 Rplp0 Eif4a2 Rps15 Actn4 Sat1 Rpl14 Hspb1 Cirbp Eif3k Rpl36al Ctsb Cfl1 Nampt Pkm Rpl18 Rhob Slc25a5 Map1lc3b Wsb1 Mrfap1 Jmjd1c |
| AR Target Genes | Concluded from reference 7 |
|  | Klk1b8 Pmepa1 Abcc4 Nkx3-1 AA986860 Fkbp5 Acsl3 Zbtb10 Herc3 Ptger4 Mphosph9 Eaf2 Med28 Nnmt Maf Gnmt Cenpn Ell2 Tmprss2 |
| Human Prostate Peripheral Zone Score | Concluded from reference 8 |
|  | Sim2 Thy1 Mycbpap Nnmt Foxa2 Bcl2a1 Apln Camk2b Map1b Cdh4 |
| AR Lineage | Lineage |
|  | Nkx3-1 Msmb Ar Tgm4 Krt18 Pla2g2a Dpp4 |
| Esr1 Target Gene | Esr1 Target Gene |
|  | mt-Nd5 Abcg1 Tiam1 Pxmp4 Btg3 Pcna Sacs Tff1 Rab31 Mcm5 Rab27b Mcm10 Dynlt3 Ret Cap2 Slc27a2 Mcm4 Cyp24a1 Cxcl12 Pole2 Adcy9 Rab26 Tpbg Nab2 Kidins220 Greb1 Recql Tsc22d3 Gla Sox3 Itpk1 Cdc25a Sgk1 Olfm1 Chrna5 Skp2 Snx24 Abat Pnrc1 Fkbp8 Ndrg4 Myof Pla2g2f Cenpu Calcr Hnrnpd Papss2 Manea Grin2b Sgk3 Sgk3 Nemp1 Ccne2 Zwilch Rrm1 Slc39a8 Rai14 Gins3 Gins2 Cdc6 Tmpo Fen1 Fhl2 Dtl Celsr2 Vamp2 Vamp2 Rfc4 Mcm6 Mcm2 Hspb8 Slc16a1 Smc4 Siah2 Med13l Abcg4 Igfbp4 Ncapg2 Prss23 Tiparp Sox13 Susd4 Esr1 |

**References**

1.Barros-Silva JD, Linn DE, Steiner I, et al. Single-Cell Analysis Identifies LY6D as a Marker Linking Castration-Resistant Prostate Luminal Cells to Prostate Progenitors and Cancer. Cell Rep 2018;25:3504-3518.e6

2.Sackmann Sala L, Boutillon F, Menara G, De Goyon-Pélard A, Leprévost M, Codzamanian J, et al. A rare castration-resistant progenitor cell population is highly enriched in Pten-null prostate tumours. J Pathol 2017;243:51-64

3.Karthaus WR, Hofree M, Choi D, et al. Regenerative potential of prostate luminal cells revealed by single-cell analysis. Science 2020;368:497-505

4.Zhang D,Park D,Zhong Y, et al. Stem cell and neurogenic gene-expression profiles link prostate basal cells to aggressive prostate cancer. Nat Commun 2016;7:10798

5.Hu WY, Hu DP, Xie L, et al. Isolation and functional interrogation of adult human prostate epithelial stem cells at single cell resolution. Stem Cell Res 2017;23:1-12

6.Henry GH, Malewska A, Joseph DB, et al. A Cellular Anatomy of the Normal Adult Human Prostate and Prostatic Urethra. Cell Rep 2018; 25:3530-3542.

7.Mulholland DJ, Tran LM, Li Y, et al. Cell autonomous role of PTEN in regulating castration-resistant prostate cancer growth. Cancer Cell 2011;19:792-804

8.Vellky JE, Wu Y, Moline D, et al. Single-cell RNA sequencing of human prostate basal epithelial cells reveals zone-specific cellular populations and gene expression signatures.J Pathol 2024;262:212-225.
