## Supplemental Table 4 for "Stem-like Prostate Remodeling in Obesity Mediates Resistance to 5α-Reductase Inhibition Therapy in BPH"

**Table S4: scRNA-seq sample information.**

| Sample ID | Genotype | Diet groups | Cell count | Median genes per cell | Mean read per cell | Number of reads | Total genes detected |
| --- | --- | --- | --- | --- | --- | --- | --- |
| M1 | *Srd5a2+/-* | RFD | 6457 | 1256 | 5871.439 | 37911884 | 24,910 |
| M2 | *Srd5a2+/-* | RFD | 9834 | 2106 | 9261.459 | 91077187 | 27,031 |
| M3 | *Srd5a2+/-* | HFD | 3829 | 2851 | 19575.308 | 74953856 | 26,330 |
| M4 | *Srd5a2+/-* | HFD | 12985 | 1016 | 5450.569 | 67790643 | 24,308 |
| M5 | *Srd5a2-/-* | RFD | 9327 | 1754 | 13048.317 | 121701655 | 26,552 |
| M6 | *Srd5a2-/-* | HFD | 3576 | 1680 | 8878.298 | 31748792 | 24,488 |
| M7 | *Srd5a2-/-* | HFD | 4597 | 2263 | 13977.139 | 64252907 | 26,148 |
